# Mitochondrial morphology provides a mechanism for energy buffering at synapses

**DOI:** 10.1101/643551

**Authors:** Guadalupe C. Garcia, Thomas M. Bartol, Sébastien Phan, Eric A. Bushong, Guy Perkins, Terrence J. Sejnowski, Mark H. Ellisman, Alexander Skupin

**Affiliations:** Luxembourg Centre for systems biomedicine, University of Luxembourg, Belvaux, L-4367; Computational Neurobiology Laboratory, Salk Institute for Biological Studies, La Jolla, CA 92037; National Center for Microscopy and Imaging Research, Center for Research in Biological Systems, University of California, San Diego, La Jolla, CA 92093

## Abstract

Mitochondria as the main energy suppliers of eukaryotic cells are highly dynamic organelles that fuse, divide and are transported along the cytoskeleton to ensure cellular energy homeostasis. While these processes are well established, substantial evidence indicates that the internal structure is also highly variable in dependence on metabolic conditions. However, a quantitative mechanistic understanding of how mitochondrial morphology affects energetic states is still elusive. To address this question, we here present an agent-based dynamic model using three-dimensional morphologies from electron microscopy tomography which considers the molecular dynamics of the main ATP production components. We apply our modeling approach to mitochondria at the synapse which is the largest energy consumer within the brain. Interestingly, comparing the spatiotemporal simulations with a corresponding space-independent approach, we find minor space dependence when the system relaxes toward equilibrium but a qualitative difference in fluctuating environments. These results suggest that internal mitochondrial morphology is not only optimized for ATP production but also provides a mechanism for energy buffering and may represent a mechanism for cellular robustness.

## 1 Introduction

Mitochondria are subcellular organelles well-known as the powerhouses of eukaryotic cells where metabolic substrates are converted to adenine triphosphate (ATP), the main energy substrate of life (Alberts et al., 2015). Dependent on their physiological context, mitochondria exhibit diverse phenotypes and their dysfunction is linked to diverse metabolic diseases and also to cancer (Wallace, 2012), diabetes (Lowell and Shulman, 2005) and neurodegeneration (Knott et al., 2008). The specific energetic needs of the brain and in particular of synaptic transmission is accompanied by a distinct mitochondrial phenotype on the molecular as well as on the morphological level (Devine and Kittler, 2018). Impairment of presynaptic homeostasis caused by mitochondrial dysfunction is believed to contribute significantly to neurodegeneration (Devine and Kittler, 2018), and compromised mitochondrial morphology is correlated with insufficient ATP production (Siegmund et al., 2018). Hence, understanding the interplay between molecular and morphological features of mitochondria may provide new insights into brain energy homeostasis and mechanisms of neurodegeneration.

The mitochondrial structure is characterized by two membranes, with one membrane surrounding the other (Figure 1A,B). The inner membrane (IM) exhibits a number of invaginations and infoldings called *cristae*. This complex structure forms specific compartments: the intercristal space (ICS), the narrower intermembrane space (IMS), and the internal matrix compartment (Figure 1C). The concrete morphology of mitochondria exhibit a large heterogeneity dependent on metabolic conditions (Hackenbrock, 1966, 1968) and is associated with distinct physiological states and their specific subcellular energy demands (Perkins and Ellisman, 2011). Within the brain, mitochondria typically exhibit a composition of lamellar and tubular cristae (Perkins et al., 2001) and synaptic mitochondria, in particular, are further specialized to their physiological context by their smaller volume (Perkins et al., 2001), higher ratio of cristae to outer membrane (OM) surface (Perkins et al., 2010) and distinct metabolic profiles (Devine and Kittler, 2018).

**Figure 1:**
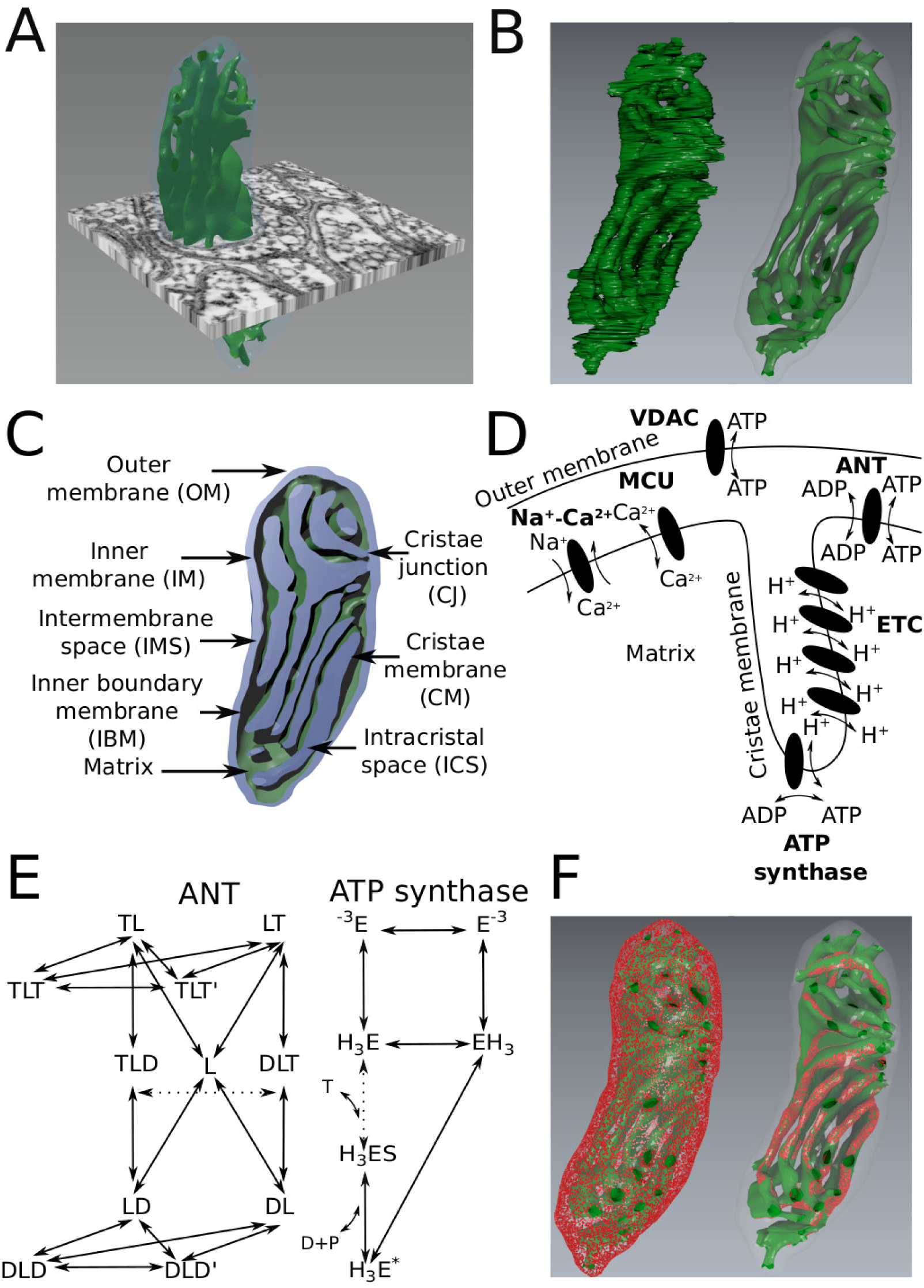
A physiological mitochondrion model based on EM tomography and dynamic simulations using MCell. (*A*) Serial electron-tomogram of a presynaptic mitochondrion in a cerebellum mouse neuron. (*B)* Morphology reconstruction first leads to mesh generation (left) and after optimization to waterproofed *in silico* representation (right) where green corresponds to the cristae membrane and semi-transparent blue membrane to the outer membrane. (*C)* Functional annotation of the reconstructed mitochondrion. The outer membrane (OM) and the inner membrane (IM) are separated by the intermembrane space (IMS). IM separates the interior matrix of the mitochondrion from the IMS and can be topologically divided into the inner boundary membrane (IBM) and the cristal membrane (CM) connected by tubular cristal junctions (CJs). (*D)* Schematic representation of main proteins involved in ATP production. The electron transport chain (ETC, not included in the model) pumps protons (H^+^) into the cristae building up a chemo-electrical gradient which is used by the ATP-synthase to phosphorylate ADP to ATP in the matrix. ATP is transported by the ATP/ADP translocator (ANT) from the matrix into the IMS/ICS and by VDAC channels into the cytosol. (*E)* Markov chain models of ANT (left) and ATP synthase (right) describing the molecular kinetics (Methods). The essential ATP-ADP exchanging step of the ANT and the ATP generating step of the synthase are indicated by the dashed arrows, respectively. (*F)* To investigate the dynamic effect of morphology in our simulations, ANTs were either homogeneously distributed on the IBM (left), co-localized with ATP synthases at the curvatures of cristae (right), or in both locations (not shown).

While extended literature (Scalettar et al., 1991; Mannella, 2006; Perkins and Ellisman, 2011) suggests a link between the inner membrane morphology and mitochondrial function, a mechanistic understanding is still lacking. This gap is mainly caused by the static data generated by electron tomography needed to reveal the internal structure. Dynamic effects caused by changing morphologies rely therefore on computational modeling. Previous simulations of the interplay between morphology and the electrochemical potential predicted an increased proton concentration in the ICS compared to the IMS (Song et al., 2013). Effects on diffusion due to the internal structure were studied based on simplified geometries (Partikian et al., 1998; Dieteren et al., 2011) and indicated anomalous diffusion in some cases (Ölveczky and Verkman, 1998), but disagreed on the impact of the internal structure (Dieteren et al., 2011; Partikian et al., 1998).

Since the interplay of diffusion with active molecules like transporters, proton pumps and synthases may have strong implications for the emergent dynamics dependent on spatial arrangement, we developed a more realistic spatiotemporal mitochondrial model (Figure 1) to (i) measure diffusion properties in a concrete physiological geometry, (ii) investigate how the interplay between diffusion and spatial localization of ATP/ADP translocator (ANT) and ATP synthase affect mitochondrial ATP production, and (iii) analyze potential energetic consequences for synaptic transmission. For this systematic investigation, our three-dimensional model is based on realistic morphologies reconstructed from electron microscopy tomograms and uses Markov state transition models to describe the molecular dynamics of ANT, ATP synthase, and the voltage-dependent anion channel (VDAC) (Figures 1D,E). We used this static geometry for dynamic simulations of the molecular interplay with spatially distinct molecular arrangements (Figure 1F). The model was implemented in MCell (Kerr et al., 2008), an agent based reaction-diffusion simulator, and compared with a corresponding space-independent ODE (ordinary differential equation) approach.

We applied our model to a synaptic mitochondrion to analyze how brain specific morphology affects ATP production capacity. Interestingly, we found that morphology has only minor effects when the system relaxes towards an equilibrium steady state condition but spatial effects are amplified in non-equilibrium situations and may provide an energy buffering mechanism in more physiologically relevant conditions of a highly dynamic environment like the synapse.

## 2 Results

### 2.1 Mitochondrial morphology reconstruction

Due to technical reasons, three-dimensional reconstructions of whole mitochondria are rare and accurate volume and surface measurements of mitochondria are lacking. We therefore initially focused on the comprehensive reconstruction of a synaptic mitochondrion from a serial electron tomogram volume (Figure 1A). The resulting reconstruction was subsequently optimized to enable dynamic simulations and detailed morphological characterization (Table 1) including the volume of 0.04 μm^3^ with a maximal length of 0.8 μm and width of 0.29 μm and 45 cristae. Based on the physiological classification of mitochondria (Figure 1C), we determined the size of the different compartments where the IMS occupies approximately a relative volume to the outer membrane of 0.27, the matrix 0.52 and the ICS 0.21.

**Table 1:**
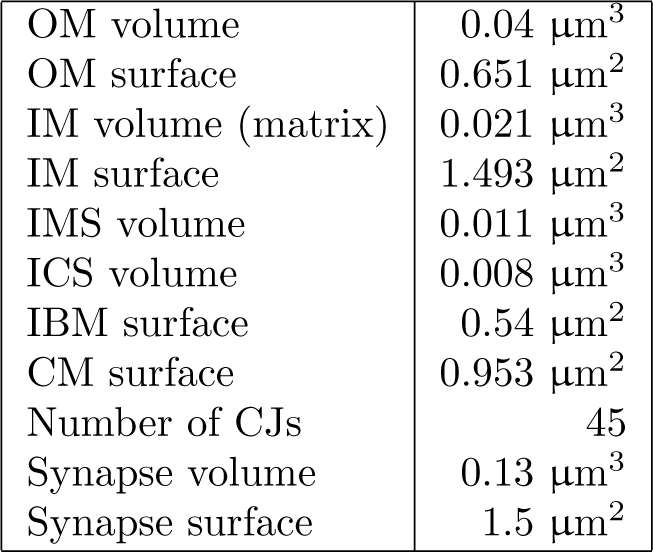
Properties measured on the reconstructed mitochondrion.

|  |  |
| --- | --- |
| OM volume | 0.04 $\mu\text{m}^3$ |
| OM surface | 0.651 $\mu\text{m}^2$ |
| IM volume (matrix) | 0.021 $\mu\text{m}^3$ |
| IM surface | 1.493 $\mu\text{m}^2$ |
| IMS volume | 0.011 $\mu\text{m}^3$ |
| ICS volume | 0.008 $\mu\text{m}^3$ |
| IBM surface | 0.54 $\mu\text{m}^2$ |
| CM surface | 0.953 $\mu\text{m}^2$ |
| Number of CJs | 45 |
| Synapse volume | 0.13 $\mu\text{m}^3$ |
| Synapse surface | 1.5 $\mu\text{m}^2$ |

### 2.2 Isolated Scenario of Equilibration

To investigate the effect of the morphology on mitochondrial dynamics, we first considered a minimal configuration and simulated only the interplay between ANT and ATP synthase dependent on their spatial arrangements (Figure 2A). In this scenario, ADP molecules corresponding to a free ADP concentration of 900 μM in the IMS and ICS (referred together as outside) are imported into the matrix by 20,000 ANT molecules and subsequently phosporylated to ATP by 3800 ATP synthases. The generated ATP can be eventually exported into cristae and the IMS by ANTs (Figure 2B-E).

**Figure 2:**
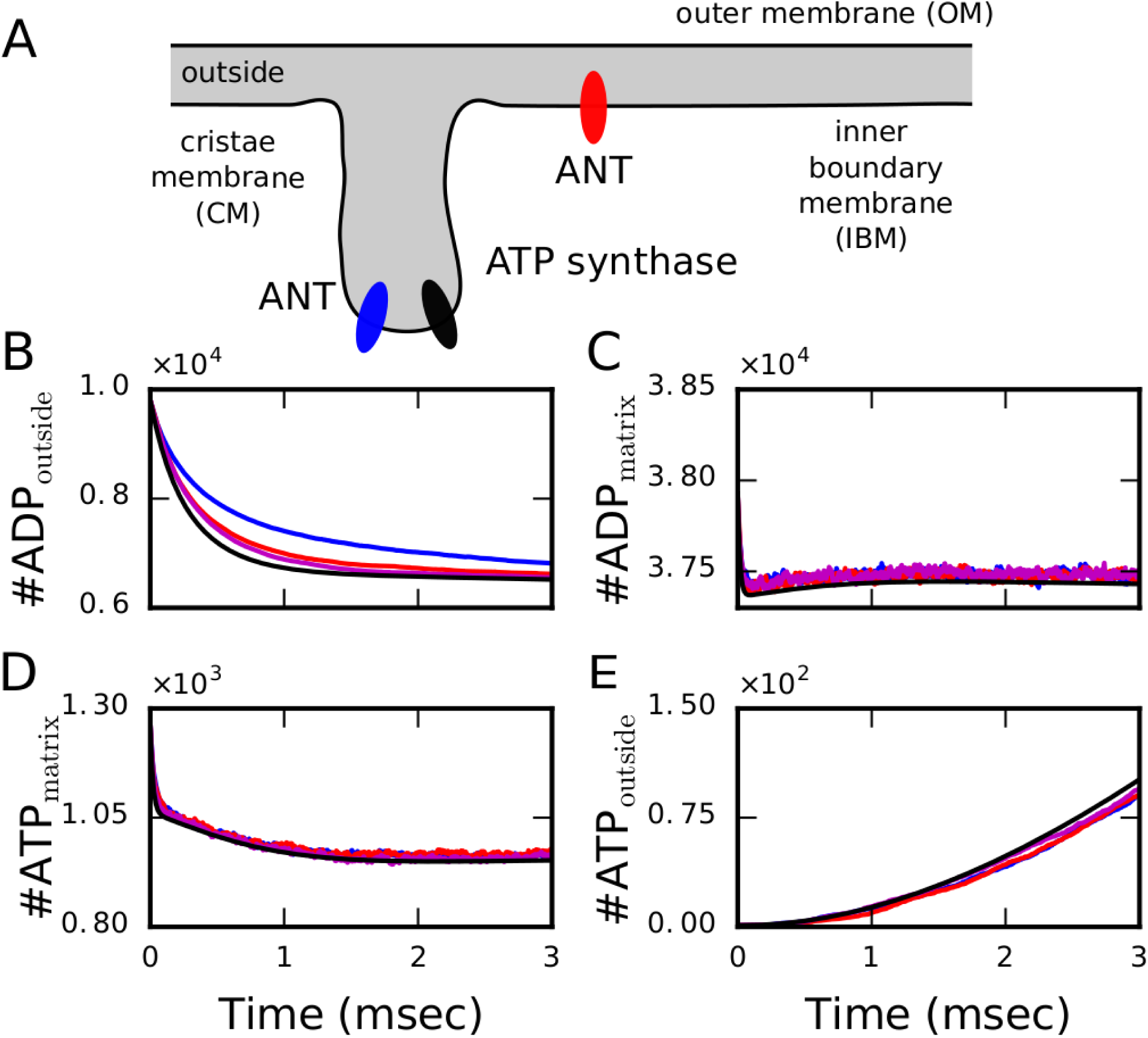
The isolated scenario of dynamic equilibration does not exhibit strong dependence on the spatial arrangement. (A) Schematic representation of the considered components and their arrangement, with ANTs either co-localized with ATP synthases in the cristae (blue), placed exclusively at the IBM (red) or in both locations (magenta). All trajectories are averaged over 10 different initial conditions and compared with the ODE system (black). (B) Simulations start with a number of ADP molecules outside, corresponding to a concentration 900 μM, which is subsequently imported into the matrix (C) and phosphorylated to ATP (D). (E) After export of ATP from the matrix, it accumulates outside. The different arrangements of ANTs do not exhibit significant differences with the corresponding ODE system.

During the equilibration process, we only observe minor differences between the different spatial arrangements within the first milliseconds which are caused by diffusion-induced delays. Nevertheless, these differences are rather small and specifically the exported ATP does neither exhibit a significant dependence on morphology nor on the molecular spatial arrangement.

### 2.3 Non-Equilibrium Induced Gradients

To investigate the mitochondrial dynamics under a more physiological non-equilibrium condition, we clamped the concentration of ADP at the surface of the OM to 900 μM, mimicking unlimited ADP resources in the cytosol, and included VDAC channels in the OM (Figure 3A) to export ATP into the cytosol. For this extended model, we monitored again the main variables of the system including the amount of exported ATP in dependence on the different spatial arrangements and compared averaged trajectories with the corresponding ODE system (Figures 3B-F).

**Figure 3:**
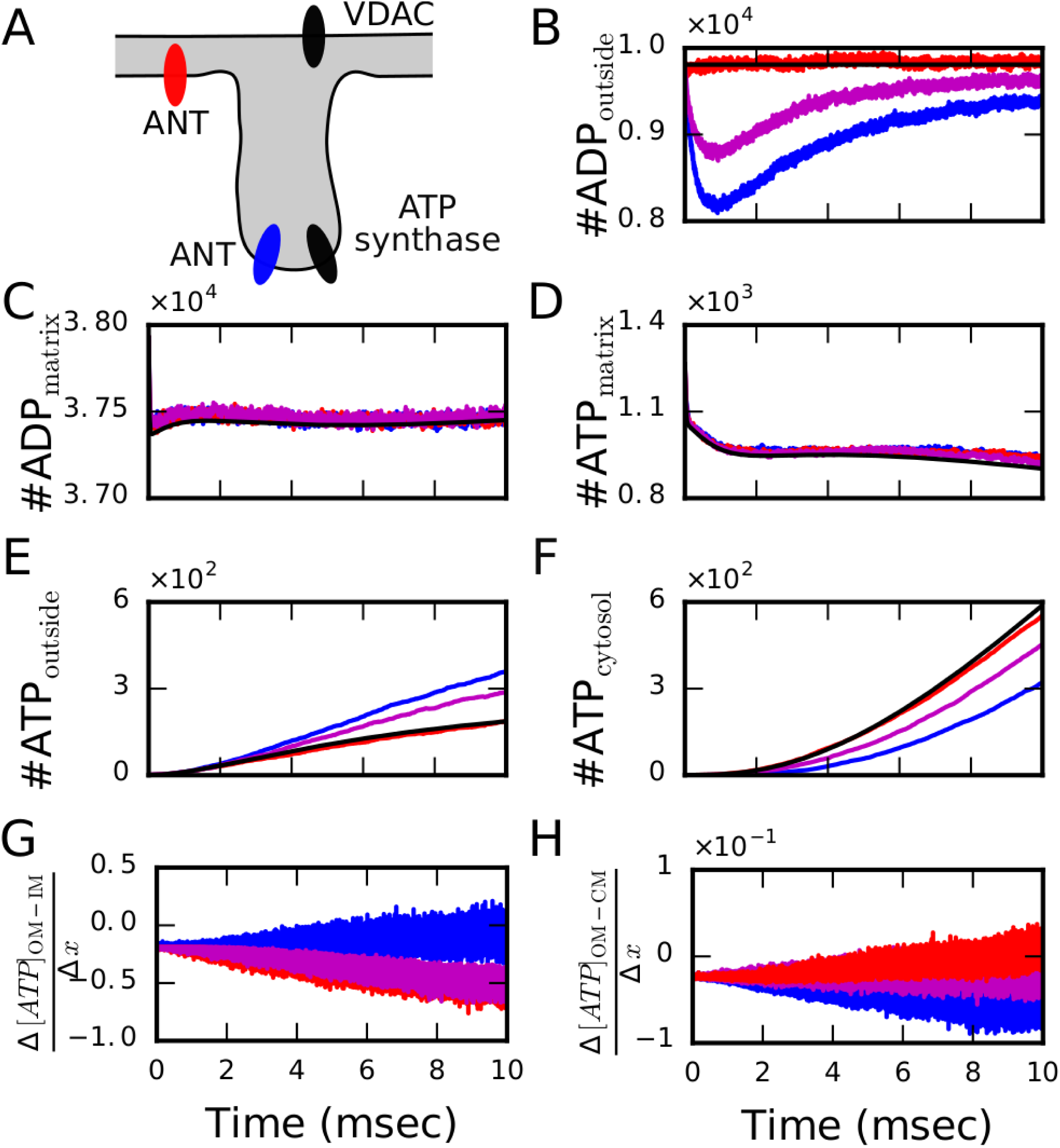
Non-equilibrium dynamics of the synaptic mitochondrion driven by clamped ADP concentration and ATP export. (*A*) Schematic representation of the components included in these simulations. (*B-H)* Comparison of averaged molecule trajectories for the distinct ANT localizations (ANTs homogeneously distributed in the IBM in red; ANTs colocalized with ATP synthase at the most curved region of the CM in blue; ANTs in both locations in magenta) with results of the ODE system (in black) exhibits most significant spatial effects for colocalization (blue) which are induced by sub-organelle ATP gradients (G,H).

In this driven system, different ANT configurations lead to distinct dynamics. When ANTs are distributed in the inner boundary membrane (IBM, red), the outside ADP concentration is almost constant but for ANTs located in the cristae membrane (CM, blue) an initial drop in the ADP concentration is caused due to a local depletion of ADP in the ICS (Figure 3B). Initially, all ADP molecules are homogeneously distributed in the outside space consisting of IMS and ICS. If ANTs are located in the IBM (red), ADP molecules are quickly bound to free ANT proteins but ADP molecules are immediately replenished from the clamped membrane concentration. Hence, no local gradients are formed. If ANTs are located in the CM exclusively (blue), ADP molecules in the ICS are quickly bound to free ANT proteins leading to a decrease of the ADP concentration in the cristae volume. This induced concentration gradient transitorily attracts more molecules from the IMS. Since the replenishment relies on slow diffusion through tubular junctions (CJs) of small diameters (∼25.5 nm in our reconstructed mesh) connecting the cristae with the peripheral volume, the drop in the outside ADP is enhanced in amplitude as well as in duration. To further characterize this scenario, we estimated the concentration dynamics in the IMS and the ICS (Supplementary Information, Figures 1B,C) and found that the initially induced ADP gradient is reducing over time and represents the driving force for the persistent differences in the outside ADP between the different configurations (Figure 3B).

The differences in the outside ADP concentrations are accompanied with differences in the outside ATP concentration (Figure 3E) where more ATP is present in the outside if ANTs are distributed in the CM (blue). In this configuration, ATP molecules are exported into the cristae volume from where they first have to diffuse into the IMS to react with VDAC proteins in the OM for export from the mitochondrion. This diffusive transport takes longer compared to the scenario when ATP is directly exported to the peripheral volume (e.g. when ANTs are located in the IBM, red). Therefore, when ANTs are in the CM, more ATP molecules are found in the outside space because they are more persistent in the ICS. To understand this interplay in more detail, we estimated the trajectories of ATP concentrations in the IMS and ICS (Supplementary Information, Figure 1F,G) and quantified the resulting gradients (Figure 3G,H, Supplementary Information, Section 1). The larger and negative ATP gradients between the OM and IBM when ANTs are located in the IBM (red) facilitate ATP transport towards the cytosol (Figure 3F) and deliver approximately double the ATP amount compared to ANTs located in the CM. Remarkably, in this non-equilibrium scenario, the setup with ANTs in the IBM does not exhibit any major differences to the space-independent ODE model whereas localization of ANTs in the cristae induces diffusion limitation for cytosolic ATP export.

### 2.4 Morphologically Buffered Energy Production at a Presynaptic Terminal

After model establishment and finding significant differences in the cytosolic ATP production in dependence on the spatial arrangement, we were interested in potential physiological consequences of morphology on the synaptic dynamics. For this purpose, we investigated the ATP production rate of the mitochondrion in its physiological context, the presynaptic terminal (Figure 4A), and included ATP-consuming reactions at the synaptic membrane to emulate the arrival of an action potential at the terminal by varying the rate constant *k*_cha_ of the ATP-consuming reactions. Based on estimations of the energetic costs of a glutamatergic synapse, we set the basal ATP consumption rate to *k*_cha_ = 2.5 · 10^4^ (Ms)^−1^ and the energy demand during an action potential to *k*_cha_ = 1 · 10^6^ (Ms)^−1^ (Supplementary Information, Section 2 for parameter estimation). To study how synaptic activation induces a transient transition between the approximated steady states for the different scenarios (Figure 4B-F), we simulated the energetic response during a 5 ms lasting recovery phase between 2 spikes by modulating *k*_cha_ as step functions between the basal and active ATP consumption rates (Figure 4G).

**Figure 4:**
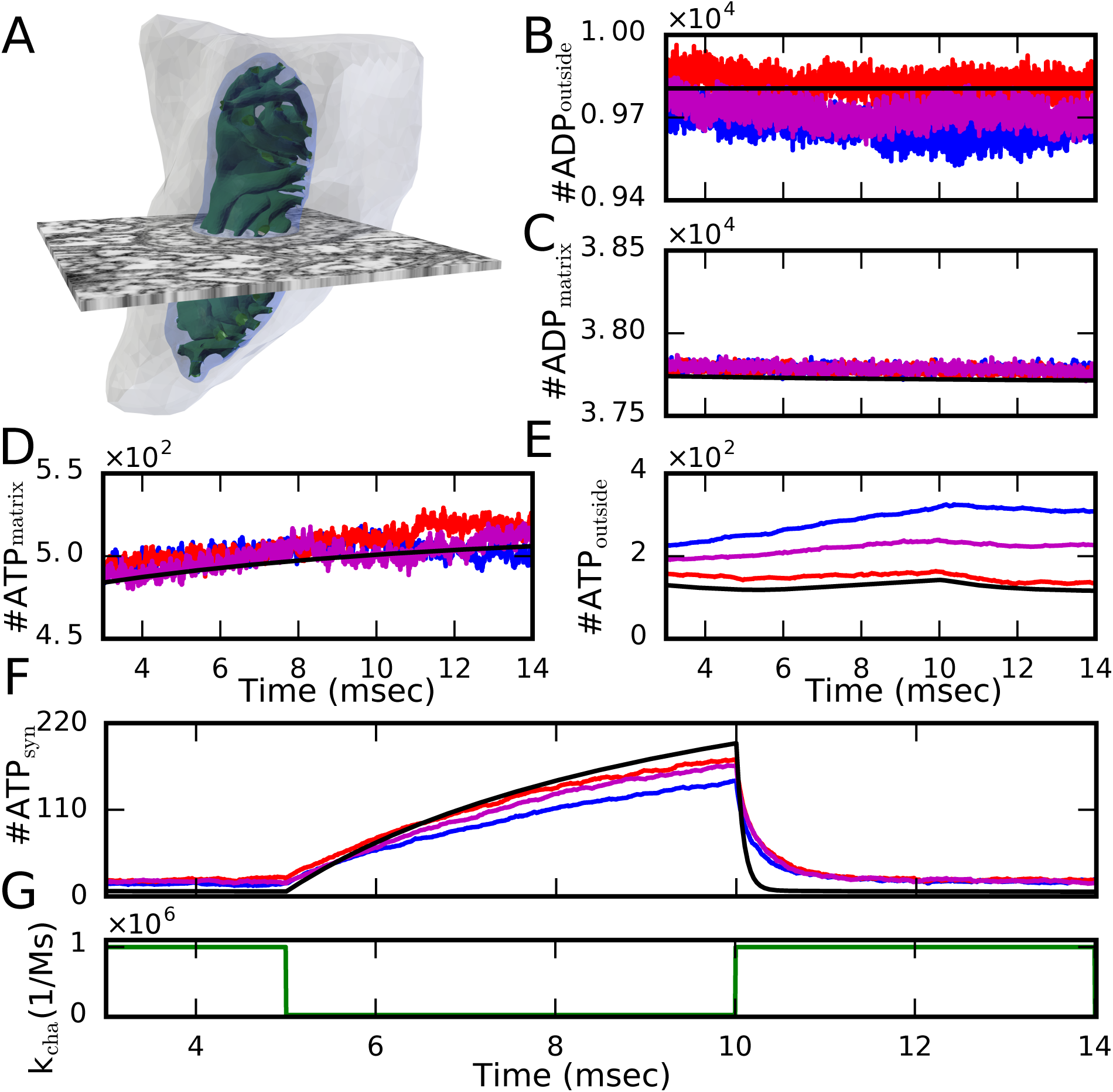
The energy dynamics at a presynaptic terminal. (*A*) In this configuration, the mitochondrion is placed within the segmented synapse (grey surface) and we considered ATP-consuming reactions. (*B-F)* Comparison of averaged molecule trajectories in the different compartments (color) and corresponding ODE results (black) for simulations of an action potential arrival at the presynaptic terminal by modulating the rate constant *k*_cha_ of the ATP-consuming reactions (*G)*. As before, the concentration of ADP in the OM was clamped and VDAC channels included in the OM. ANT molecules were placed in three different locations: ANTs homogeneously distributed in the IBM (red), ANTs colocalized with ATP synthase at the most curved region of the CM (blue), and ANTs in both locations (magenta).

In contrast to the previous scenario, these simulations started close to steady state conditions determined by the ODE system. Therefore, we did not observe an initial dip in the outside ADP concentration (Figure 4B) but a stable gradient that drives the differences among the distinct configurations. Due to ADP clamping, the outside ADP concentration stays constant for the ODE approach and similarly for the scenario where ANTs are localized in the IBM (black and red in Figure 4B, respectively), whereas for ANTs exclusively or partly localized in the cristae, a small drop is observed (blue and magenta in Figure 4B, respectively). Interestingly, ADP as well as ATP concentrations in the matrix are slightly increased consistently for all spatial simulation compared to the ODE approach (Figure 4C-E).

The most predominant difference is subsequently observed in the outside ATP concentration where localization of ANTs in the IBM again exhibit similar concentrations as the ODE system whereas localization of ANTs in the cristae lead to substantially increased ATP levels (red and blue in Figure 4E, respectively). As in the non-equilibrium scenario, this increase is caused by ATP within the ICS from where it first has to diffuse to the IMS for subsequent export into the cytosol. Hence, the cytosolic ATP is slightly lower for ANTs located in the cristae compared to ANTs in the IBM (blue and red in Figure 4F). Despite this difference, all spatiotemporal scenarios exhibited consistently smaller ATP amounts within the synapse during the recovery period with low energy demand compared to the ODE simulations (24% less for co-localization vs 10% less for IBM localization). After synaptic activation, the spatiotemporal simulations displayed a slower decrease in synaptic ATP and reduced differences in the ATP concentration between base level and activation conditions (relative change of 0.87 for co-localization vs 0.88 for IBM localization vs 0.97 for ODE). These results together indicate that ATP molecules can be buffered by the complex morphology and support adaptation to variable conditions.

We finally used the presynaptic model to calculate net ATP production rates from the second peak in Figure 4F. For ANTs located in the IBM we calculate a rate of ∼ 31 molecules/ms slightly reduced compared to the ODE system (∼ 38 molecules/ms). The model with ANTs exclusively in the CM exhibits a rate of ∼ 26 molecules/ms. Comparison with theoretical estimations (Table 2) and approximations in the literature exhibit good agreement.

**Table 2:**
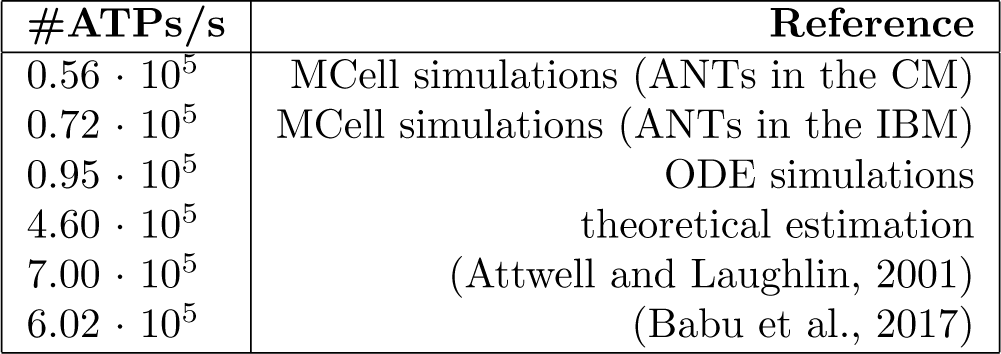
Estimation of ATP production in synaptic mitochondria.

| **#ATPs/s** | **Reference** |
| --- | --- |
| $0.56 \cdot 10^5$ | MCell simulations (ANTs in the CM) |
| $0.72 \cdot 10^5$ | MCell simulations (ANTs in the IBM) |
| $0.95 \cdot 10^5$ | ODE simulations |
| $4.60 \cdot 10^5$ | theoretical estimation |
| $7.00 \cdot 10^5$ | (Attwell and Laughlin, 2001) |
| $6.02 \cdot 10^5$ | (Babu et al., 2017) |

## 3 Discussion

Mitochondrial morphology is thought to be context dependent and a mechanism to adapt to specific energetic requirements (Scalettar et al., 1991; Mannella, 2006). Mitochondria in the brain and specifically at synapses exhibit rather unique and complex morphologies (Perkins et al., 2001, 2010) that may reflect the high energy demand for neuronal information transmission (Attwell and Laughlin, 2001). Since the internal structure of mitochondria can be only resolved by EM tomography, a mechanistic understanding of how morphology is affecting mitochondrial dynamics relies on mathematical modeling to simulate dynamic consequences from the static images.

While modeling approaches have estimated the morphological effect on the mitochondrial membrane potential (Song et al., 2013) and diffusion properties based on simplified geometries (Dieteren et al., 2011; Ölveczky and Verkman, 1998), the consequences for the main function of ATP production of a real physiological morphology is only vaguely understood. Here, we used an electron tomogram of a presynaptic terminal in mouse cerebellum to (i) comprehensively reconstruct and analyze in detail the morphology of an entire mitochondrion (Table 1) and to (ii) subsequently investigate the dynamic consequences of the interplay between the complex morphology the spatial molecular orchestration by our developed computational model based on the mitochondrial morphology and molecular properties of the main adenosine phosphate processing molecules.

Surprisingly, simulations of the isolated scenario without any ADP import from and ATP export into the cytosol do not exhibit a strong dependence on the spatial arrangement (Figure 2) indicating that the assumed diffusion properties do not lead to a diffusion limiting condition. In accordance with theoretical considerations, comparing the timescales of diffusion and reactions indicated only a slight overlap for this scenario (Supplementary Information, Section 2). A morphological effect on ATP production could only be found for diffusion coefficients decreased by two orders of magnitude (Supplementary Information, Section 1). Although some studies (Scalettar et al., 1991; López-Beltrán et al., 1996; Dieteren et al., 2011) showed evidence of severe hindrance of diffusion in the matrix, more recent experiments estimated that diffusion is only three to four fold smaller than in water (Partikian et al., 1998). In our model, we reduced the diffusion coefficient of ATP and ADP by one order of magnitude to reflect their ionized form and related interactions with other charged particles. The potential interaction of the ions with the membrane potential leading to electro-diffusion are not included in the current model but could actually decrease diffusion further and induce a regime of diffusion limitation. Independent of the diffusion limitation, our simulations indicated anomalous diffusion in agreement with previous evidences (Ölveczky and Verkman, 1998).

Although diffusion had only a minor effect in the isolated system, spatial aspects became significant when bringing the mitochondrion in contact with the cytosol under unlimited access to ADP and ATP export through VDAC (Figure 3). Under these more physiological conditions, the spatial organization of ANTs had a significant effect on ATP gain within the cytosol. While the spatiotemporal simulations did not exhibit significantly strong deviations from the spatially independent ODE system when ANTs were exclusively located at the IBM, the co-localization of ANTs with ATP synthases at the apex of cristae led to an approximately halved ATP export into the cytosol. Careful analysis of the dynamics revealed that this effect is driven by smaller concentration gradients between the ICS and the OM for ANT localization in the cristae, which led to ATP buffering within the cristae. This scenario is in contrast with the greater concentration gradient formed between the IBM and the OM when ANTs are located in the IBM what is facilitating ATP transport into the cytosol. These findings quantitatively support the importance of sub-organelle gradients suggested in the literature (Mannella, 2000). To test whether this buffering mechanism might have an effect on synaptic physiology and explain the distinct morphology of brain and specifically of synaptic mitochondria, we subsequently simulated the mitochondrion in its synaptic environment with a variable cytosolic ATP consumption reflecting changes during synaptic transmission. These simulations have shown that ATP buffering in cristae caused by the non-equilibrium induced gradients is a mechanism to buffer large energy demand peaks.

We finally used our detailed model to calculate the ATP production rate of the considered mitochondrion for the different scenarios. The resulting rates of ∼10^5^ molecules of ATP per second are in agreement with our theoretical estimation based on the ANT translocation rate and the ANT density in mitochondria (Supplementary Information, Section 2). These values are further supported by independent approximations found in the literature (Babu et al., 2017; Attwell and Laughlin, 2001) as summarized in Table 2 where minor deviations to the calculations by Attwell and Laughlin (2001) would rematch for firing rates of 30 Hz. The main mechanism how mitochondria decode the firing rate is probably Ca^2+^ influx through the mitochondrial calcium uniporter (MCU) (Kirichok et al., 2004). Incorporating the MCU and the effect of Ca^2+^ on the membrane potential in a future version of the model will allow for more detailed predictions of ATP production rates in dependence on neuronal activity.

Overall, our systematic approach with our detailed mitochondrial model has shown that the concrete morphology of the presynaptic mitochondrion induces anomalous diffusion but has not per se an impact on ATP production when the system relaxes towards an equilibrium steady state (Figure 2). In contrast, the spatial arrangement of ANTs under non-equilibrium conditions induce sub-organelle gradients that led to a significant effect on the cytosolic ATP concentration (Figure 3F). Physiological simulations of the synaptic dynamics suggest that this buffering effect might be a mechanism to smear out the variable energy demands (Figure 4) and may therefore increase robustness and adaptability of synapses and explain the distinct morphology of brain mitochondria.

## 4 Material and Methods

Spatiotemporal simulations were performed with MCell (version 3.4) (Kerr et al., 2008) and compared with space-independent simulations of the corresponding rate equation system. For the spatiotemporal model, each molecular component was first implemented independently, parameterized and validated by experimental data and eventually combined in the realistic mitochondrial model. The entire dynamical system has 21 variables (6 for ATP synthase, 11 for ANT and 4 for ADP and ATP concentrations within the 2 compartments).

### 4.1 Specimen Preparation

A 1-month old C57BL/6NHsd male mouse was anesthetized with ketamine / xylazine and transcardially perfused with Ringer’s solution followed by 2.5% glutaraldehyde, 2% formaldehyde, 2 mM CaCl_2_ in 0.15 M sodium cacodylate buffer. The fixation was started at 37°C and the fixative was cooled on ice during perfusion. The brain was post-fixed after removal from the cranium in the same fixative solution for 1 hour at 4°C. The cerebellar vermis was cut into 100 μm thick sagittal slices on a vibrating microtome in ice-cold 0.15 M cacodylate buffer containing 2 mM CaCl_2_ and briefly stored in same buffer prior to HPF. A 1.2 mm tissue punch was taken from a tissue slice and placed into a 100 μm deep membrane carrier filled with 20% bovine serum albumin in cacodylate buffer and frozen with an EM PACT2 HPF apparatus. The specimen was freeze substituted in extra dry acetone (Acros) using an AFS2 as follows: 0.1% tannic acid at −90°C for 24 hours, wash 3x 20 min in acetone, 2% OsO_4_ / 0.1% uranyl acetate at −90°C for 48 hours, warmed for 15 hours to −60°C, held at −60°C for 10 hours, and warmed to 0°C over 16 hours. The specimen was infiltrated with a series of Durcupan ACM: acetone solutions and then embedded in 100% Durcupan at 60°C for 48 hours.

### 4.2 Electron Tomography

300 nm sections were cut and collected on 50 nm thick Luxel slot grids. The sections were glow discharged and coated with 10 nm colloidal gold. Tilt series were collected on an FEI Titan 300 kV microscope with a 4k × 4k CCD detector (Gatan Ultrascan). Four tilt series were collected from the region of interest at 0, 45, 90, and 135 degrees rotation of the specimen plane. Each tilt series was collected from −60 to +60 degrees with 1° increments. Projection images were collected with a pixel size of 0.4 nm, and images were binned by 4 prior to tomographic reconstruction with TxBR (Phan et al., 2016).

### 4.3 Model Geometry

Mitochondrial and synaptic 3D *in silico* reconstructions were performed from 3 sections of serial electron tomogram of a high pressure frozen / freeze substituted (Sosinsky et al., 2008) cerebellum sample, exhibiting final pixel resolution of 1.64 nm, leading to a stack of 360 images containing the mitochondrion and the synapse. First, membranes of the presynaptic mitochondrion were manually traced using RECONSTRUCT. Afterwards, contours were converted into three-dimensional surfaces by VolRover. Finally, meshes were imported into Blender to generate a triangulated, watertight and manifold mesh using CellBlender’s Mesh Analysis tool. Further optimization was performed with the mesh improvement library and Blender add on GAMer (Figure 1B left). To consider possible compression effects vesicles were traced, and its shape was set to spheres of diameter 40 nm. We found shrinkage in the Z direction of 20%, in order to correct for this we rescaled the reconstructed meshes by a factor of 1.239 in the Z direction. Two movies have been generated to visualize the complex morphology (Supplementary Information, Section 1).

### 4.4 Molecular ATP/ADP translocator (ANT) Model

The ANT model is based on the work of Metelkin et al. (2006). Two additional states were added to track futile translocations in MCell. The resulting kinetic ANT model (Figure 1E) is composed of 11 states and 19 bidirectional transitions between them resembling the binding and unbinding of ATP and ADP from different sides of the IM. Starting from fitted flux parameters (Metelkin et al., 2006) for ANT extracted from heart mitochondria (Kraemer and Klingenberg, 1982), we first estimated parameters for the implementation in MCell and the corresponding ODE model (Supplementary Information, Section 2). With this set of parameters, we qualitatively reproduced the independent data from published work (Kraemer and Klingenberg, 1982; Duyckaerts et al., 1980). To obtain a reference ATP turnover rate, we used published data for synaptic mitochondria (Chinopoulos et al., 2009). The complete list of parameters are given in the Supplementary Information, Table 1. The location of ANTs in mitochondria has not yet been definitively determined. Experimental evidence show on the one hand that they may form complexes with ATP synthases and phosphate carriers (Ko et al., 2003) located in the CM (Wittig and Schägger, 2009; Vogel et al., 2006) and, on the other hand, studies report an association with VDAC located in the IBM (Vyssokikh et al., 2001). In our simulations we explored the functional implications of these different locations by placing them (i) homogeneously distributed in the IBM (Figure 1F, on the left), (ii) colocalized with ATP synthase in the CM (Figure 1F, on the right) or (iii) in both locations.

### 4.5 Molecular ATP Synthase Model

The ATP synthase model is based on the six state model of a proton pump by Pietrobon and Caplan (Pietrobon and Caplan, 1985) shown in Figure 1E. A clockwise cycle starting in *E*^−3^ represents the binding of 3 protons from the IMS, transport of the protons, binding of ADP and phosphate (P_*i*_) and subsequent synthesis of ATP, followed by unbinding of the protons in the matrix. In our model, we considered the proton concentration inside the ICS as well as proton and phosphate concentrations in the matrix to be constant and used ADP and ATP in the matrix and the IMS as input variables. In our model, ATP synthases were localized at the apex of the CM in lamellar cristae and along the length of tubular cristae, in accordance to experimental findings (Strauss et al., 2008). All model parameters are given in the Supplementary Information, Table 2.

### 4.6 Molecular VDAC Model

To consider processes that exports ATP from the mitochondrion into the cytosol, we included VDACs, the main mechanism for metabolites to cross the OM. We implemented a rather basic model of VDAC assuming that VDAC proteins interact with ATP and translocate it to the cytosol by the reaction VDAC+ATP_mito_ ⇌VDAC+ ATP_cyto_. In our simulations VDAC proteins were homogeneously distributed within the OM (Supplementary Information, Section 2 and Table 3 for details and parameters values).

### 4.7 Metabolite diffusive properties and buffers

Diffusion coefficients were estimated previously based on measurements of green fluorescent protein (GFP) in the matrix of mitochondria of diverse cells (Partikian et al., 1998; Dieteren et al., 2011) reporting that the free diffusion is two to fourfold reduced compared to water (Partikian et al., 1998; Dieteren et al., 2011). For our simulations, the free diffusion coefficient is relevant since the effect of morphology is included in our model. Although GFP as a protein has a higher molecular weight than ATP or ADP and as such would have a lower diffusion coefficient, ATP and ADP are ionized in neutral solutions as ATP^4−^ and ADP^3−^ leading to lower mobility due to interactions with other charged particles and the electrochemical gradient at the membrane. To account for these interactions, we reduced the free diffusion coefficient by one order of magnitude to 1.5 · 10^−7^cm^2^s^−1^.

ADP and ATP can react with different cations, be bound or ionized. Therefore, the total concentration of ATP can be distributed in different compounds or states like ATP^4−^, ATPMg^2−^. These distributions can be estimated by coefficients representing the fraction of unbound ATP in the matrix of mitochondria or the external compartments. For our model, mitochondrial ADP^3−^ and ATP^4−^ concentrations were estimated analogously to published data (Magnus and Keizer, 1997) as [ADP]_m,free_ = 0.8 [ADP]_m_, [ATP]_m,free_=[ATP]_m_, [ATP^4−^] = 0.05 [ATP] and [ADP^−3^] = 0.45 [ADP]_free_. The concentrations of ATP and ADP in the matrix were set to 2 mM and 10 mM, respectively, and to 0.01 mM and 2 mM in the cytosol.

### 4.8 Space-independent ODE approach

For each molecular model, we also developed a corresponding ODE approach describing the fluxes based on mass action kinetics (Supplementary Information, Section 2). The ODEs were integrated by PyDSTool (Clewley et al., 2007). To investigate morphological effects, the different spatial configurations simulated with MCell were compared with corresponding solutions of the ODE system.

### 4.9 Numerical Experiments

For model establishment, we performed 3 distinct *in silico* experiments to disentangle the contribution of the different molecular components to the dynamics. In a first set of simulations, we started with a fixed number of ADP molecules and let them be phosphorylated to ATP without any export or consumption of ATP. Hence, in this *isolated scenario*, ATP molecules accumulate in the mitochondrion. In a second configuration, we consider the mitochondrion to be embedded in a cube of dimension 0.45 μm^3^ reflecting the cytosol with unlimited resources of ADP by clamping the concentration of ADP in the OM, and include VDAC channels in the OM for mitochondrial export. The more *physiological scenario* of a fluctuating energy demand at a synapse is similar to the scenario of unlimited resources but with the mitochondrion located in the reconstructed synapse. ATP-consuming reactions are included at the synaptic membrane representing different ATP-consuming processes. To reflect the activation of the reactions due to action potential arrivals we increase the rate constant of the reactions. For each configuration, we ran 10 individual simulations with different random seeds. Averaged trajectories were compared between configurations and compared with the spatially independent scenario described by the corresponding ODE sytem.

## 4.10 Acknowledgements

This work was supported by the National Research Fund of Luxembourg in the frame of a PhD Grant No.9984574 to G.C.G. and the National Institute of Health grants P41GM103412 and R01DA038896 to M.E. and P41 GM103426 to A.S.. We thank Emily Liu for her help tracing the membranes.

