## Supplementary Information for "Mitochondrial morphology provides a mechanism for energy buffering at synapses"

May 20, 2019

#### 1 Additional results

##### 1.1 Mitochondrion and synapse properties

The serial electron tomogram used to reconstruct the mitochondrion is shown in Movie 1. The resulted reconstruction is shown in Movie 2 to appreciate the complex internal structure of synaptic mitochondria. In Movie 2, the outer membrane (OM) is shown in white and after few seconds disappear, the inner membrane (IM) is transparent blue and the cristae membrane (CM) is colored in green, similar to Movie 3. In Movie 3, we also show the reconstructed synapse, and an artificial digital slice of the tomogram (the thickness is artificial), we also included ATP molecules and ANTs (ADP/ATP translocators) in the cristae membrane. This movie corresponds to a real simulation, with the same conditions as in the last experiment with the synapse, it corresponds to the first millisecond of the simulations (the time step of the visual representation is 5 μs).

The material that comes with this study is available at this doi: www.doi.org/10.17881/lcsb.20190507.01

**Movie 1.** Screen trough the serial electron tomogram of the reconstructed mitochondrion. The 3 sections of the serial electron tomogram of a high pressure frozen / freeze substituted mouse cerebellum sample exhibited a final pixel resolution of 1.64 nm and contained the complete mitochondrion in a stack of 360 images.

**Movie 2.** Reconstructed mitochondrial morphology. The outer membrane is initially shown in grey and subsequently removed to visualize the inner membrane (blue) and the cristae membrane (green) forming laminar and tubular cristae.

**Movie 3.** Exemplified mitochondrial simulations. The reconstructed mitochondrion was used to perform physiological simulations within the presynaptic terminal. The simulation visualizations shows the ANT (red) localizations in the cristae membrane (green) and the resulting ATP molecules (yellow) for 2 ms.

#### 1.2 Local concentrations and gradients

For the scenario in which the ADP concentration is clamped at the OM, we estimated local ADP and ATP concentrations in the ICS, the inner boundary (IBM) and outer membrane space for the 3 different spatial arrangements (Figure 1). For this purpose, we counted with MCell the numbers of hits in an open region from which the concentrations can be estimated by assigning a small volume to this surface.

As explained in the main text, an initial drop in the ADP concentration is observed within the ICS when ANTs (adenine nucleotide translocators) are located in the CM as shown by the blue line in Figure 1C. The opposite holds for the ATP concentration which is persistently higher in the ICS due to the constant transport of ATP from the matrix to the cristae space (blue line in Figure 1G). We computed local concentration gradients between the inner boundary and outer membrane as well as between the cristae and the outer membrane (Figure 2). To calculate these gradients, we measured the distance between the outer and inner membrane ($\leq 0.02$ μm[1]) and between the middle of the mitochondrion to the outer membrane ($\sim 0.15$ μm), respectively, by the Blender add-on Measureit.

From the resulting data, we identified a strong gradient formed between the IBM and OM if ANTs are located in the IBM (grey line in Figure 2C) driving ATP export from the mitochondrion. In contrast, ATP import is observed in case ANTs are localized in the CM by an on average positive gradient (blue line in Figure 2C). In this case with ANTs localized in the CM, the gradient between the CM and the OM points to the exterior but is one order of magnitude smaller than the gradient between the IBM and OM. These observations underlay the results obtained in all different configurations.

#### 1.3 Additional variables

Here, we include all variables of the model that are not shown in the main text for both, the averaged traces of MCell simulations and the corresponding solution of the ODE system. Figure 3A-H exhibits all variables of the ANT model for the isolated scenario of equilibration and panels I-N show all variables of the ATP synthase model. Analogously, Figure 4 exhibits the variables for the non-equilibrium condition induced by a clamped ADP concentration at the surface of the OM where Figure 4A-H shows variables of the ANT model and panels I-N of the ATP synthase model. Figure **??** shows the response of the system in the third scenario considering a symmetric function for the ATP-consuming reactions.

#### 1.4 Reduced diffusion

ATP and ADP molecules are present in an ionized form in neutral solutions what may induce an additional reduction of diffusion. Here, we explore the effect of a smaller diffusion coefficient on the dynamics. We performed analogous simulations for the 3 different scenarios described in the main text but with a one order of magnitude smaller diffusion coefficient of $D = 1.5 \cdot 10^{-8}$ $cm^2s^{-1}$. For this diffusion coefficient, substantial differences are observed in almost all cases (Figures 6, 7 and 8). Remarkably, significant less ATP molecules (only $\sim 47$) reached the cytosol after 10 ms when ANTs were located in the CM (blue line in Figure 7E). When ANTs were located in the IBM, $\sim 354$ ATP molecules reached the exterior during the same period of time (grey line in Figure 7E). This diffusion limitation based effect was also found for faster diffusion as used in the main text but reduced diffusion is amplifying the differences between configurations.

Based on these simulations, we could estimate the rate at which ATP becomes available within the cytosol what is severely delayed under diffusion limitation conditions (Figure 8). For ANTs placed in the IBM, the ATP rate is 38 molecules per millisecond, and for ANTs in the CM the rate reduces to 11 molecules per millisecond whereas the rate for the spatially independent ODE yields 62 molecules per millisecond. If diffusion is reduced, the location of ANTs have a tremendous impact on the rate at which ATP can reach the cytosol and thus on the rate of energy supply at the synapse.

[1]The maximal distance between the IBM and the OM is 0.02 μm but membranes also get closer to each other. Hence, estimated values of gradients represent lower bounds.

#### 1.5 Anomalous Diffusion

The extreme mitochondrial morphology has evoked a scientific discussion on diffusion properties within the organelle. Combining mathematical models and experimental data, the diffusive properties of proteins in the mitochondrial matrix have been measured (Partikian et al., 1998; Dieteren et al., 2011) and simulations of diffusion in obstructed objects suggested anomalous diffusion within organelles (Ölveczky and Verkman, 1998). To test this hypothesis, we used here our physiologically realistic reconstruction and simulated particle diffusion within the interior of the mitochondrion with and without the consideration of the internal structure (cristae membrane). From the resulting particle trajectories, we computed the mean-square displacement (MSD) of the molecules in both configurations. For normal diffusion, the MSD obeys Fick's law and equals $6\,\mathrm{D}\,\mathrm{t}$ where D is the diffusion coefficient (set to $\mathrm{D} = 1.5 \cdot 10^{-7}\ \mathrm{cm^2 s^{-1}}$) and t represents time. In particular, this relation predicts a linear dependence of the MSD on time t.

We compared this theoretical prediction with the MSD measured in simulations with and without internal structure (Figure 9). For the comparison, we calculated the linear regressions of the measured MSD in both configurations and determined the slopes. The slope of the theoretical prediction for Fick's law in an open space is $9 \cdot 10^{-7}\ \mathrm{cm^2 s^{-1}}$ whereas we measured $4.65 \cdot 10^{-7}\ \mathrm{cm^2 s^{-1}}$ and $4.07 \cdot 10^{-7}\ \mathrm{cm^2 s^{-1}}$ for simulations without and with considering the internal structure, respectively. While the deviation on the large time scale is caused by the closed space of the mitochondrion, the deviation on the shorter time scale is influenced by the internal structure. Therefore, the deviation of the slope from the theoretical prediction is larger and rather instantaneous for the simulations considering the internal structure whereas the simulations only considering the closed space of the mitochondrion initially exhibits a rather similar slope and only deviates from the prediction on the longer time scale.

### 2 Material and Methods

Here, we give the details on the modeling methods. For this purpose, we first introduce the molecular Markov chain models (MCMs) used to describe the ANT, the ATP synthase and the voltage dependent anion channel (VDAC) including explanations of model parameterization by experimental data. Subsequently, we describe the derivation of the corresponding ODE systems, which were used to validate the MCMs and to compare the spatiotemporal simulations with the space independent approach. Finally, we detail the estimation of the ATP production in the synapse and the time scale analysis.

#### 2.1 Markov chain models

##### ADP/ATP translocator (ANT)

The ANT model is based on published work (Metelkin et al., 2006) and it is composed of eleven kinetics states (Figure 10) that represent the protein complex L in its different binding configurations. ATP and ADP correspond to T and D, respectively, and the index i refers to the matrix (interior) side, and o to IMS and ICS that represents together the outside space. Hence, L represents the free protein and YLX describes the molecular state with one X molecule bound from the matrix side and one Y molecule bound from the outside space. The kinetic rate constants are inferred from independent experimental data (in the following Section) and given in Table 1.

The dissociation constants of the ANT exhibit a complex dependence on the membrane potential (Klingenberg, 2008). Metelkin et al. (2006) developed a model introducing the dependence into the affinity of the adenylates and the rate constants. Our model is based on this previous work. Nevertheless more parameters needed to be estimated, and therefore others had to be slightly modified. The resulting parameters slightly deviate from those used by Metelkin et al. (2006) and are given in Table 1. The dissociation constant $\mathrm{K_{T_o}}$ is changed from $\sim$ 400 μM to 500 μM in our simulations based on the parameter inference. Similarly, $\mathrm{K_{D_o}}$ changed from $\sim$ 51 μM to 25 μM . We also estimated the dissociation constants of ADP and ATP from the matrix side $\mathrm{K_{T_i}}$ leading to a value of 6.25 mM and $\mathrm{K_{D_i}}$ of 10 mM, respectively. Interestingly, these values for the matrix side are in the mM range, whereas the rates for the external side are in the $\mu$M range indicating a lower affinity of substrates on the matrix side. Similar values have been observed for the Michaelis-Menten constant Km for other mitochondrial carriers (Klingenberg, 2008).

From these dissociation constants, the backward and forward rate constants were estimated. The forward rate constants were set to be smaller than the diffusion limited rate (Milo and Phillips, 2015), and the backward rates were set to satisfy the dissociation constant. With these modifications, we were able to qualitatively reproduce the data from Kraemer and Klingenberg (1982); Duyckaerts et al. (1980) (data not shown). Although, the data was qualitatively well reproduced, the resulting turnover rate of 0.1 $\mathrm{s^{-1}}$ is two orders of magnitude smaller than the measured rate (Chinopoulos et al., 2009). This discrepancy might be caused by the limitations of the experimental data in Kraemer and Klingenberg (1982) that was generated with liposomes, and it is known that the lipid composition can affect the velocity of the translocator. To compensate for these limitations and to better calibrate our model, we included a third experiment (Chinopoulos et al., 2009) in which the transient appearance of ATP in the medium was measured for isolated synaptic mitochondria. From this data we were able to adjust the rate of ATP translocation ($k_p$) to match the measured turnover rate of 23 $s^{-1}$ (Chinopoulos et al., 2009) for synaptic mitochondria. This led to a rate constant of 800 $s^{-1}$ for the ATP exporting reaction DLT $\xrightarrow{k_p}$ TLD. Similar values have also been measured in transient experiments with liposomes (Gropp et al., 1999). All the final parameters of the ANT model used for the presented simulations are summarized in Table 1.

##### ATP synthase

The ATP synthase model is composed of six states (Figure 11A). Binding events from the matrix to the synthase E are indicated on the left and binding from the outside on the right, respectively. A clockwise cycle of the synthase starting from $E^{-n}$ corresponds to the binding of $n$ protons from the outside and transport to the matrix followed by the binding of ADP and $P_i$ for the successive synthesis of ATP and unbinding of the $n$ protons in the matrix side. The ATP generating step is indicated by the dashed arrow. The states are assigned numbers (Figure 11B), and the first order or pseudo first order rate constant of the specific transition, from state i to state j is $k_{ij}$ where we follow the same notation as in the original paper (Pietrobon and Caplan, 1985). State $^{-3}E$ corresponds to state number 1, follow by state number 2 $H_3E$, etc, following a counterclockwise rotation. All parameters values are given in Table 2.

##### ATP consuming reactions

To set the parameter values of the ATP consuming reactions, we used the calculations by Attwell and Laughlin (2001) for the energy demand of glutamatergic signaling. From these considerations, they estimated a demand of 11,000 ATP molecules per released vesicle, 12,000 ATP molecules for related $Ca^{2+}$ removal and $\sim$ 821 ATP molecules for endocytosis and exocytosis of vesicles. From the 11,000 ATP molecules needed for glutamate recycling, we only considered 1,333 (for the packing of glutamate in vesicles) since this is actually processed at the presynaptic terminal whereas the other processes are located within astrocytes. Thus, $\sim$ 14,000 ATP molecules are needed per vesicle released. For a firing frequency of 200 Hz and a release probability of 0.25, the total number of ATP molecules needed at the synapse equals $7 \cdot 10^5$ ATPs/s ($0.25 \times 200\text{Hz} \times 1.4 \cdot 10^4$).

To consider ATP consumption within the synapse, we placed channel molecules at the synaptic membrane which consume ATP by the reaction ATP + channels $\xrightarrow{k_{cha}}$ channels. For mimicking the action potential dependent ATP consumption, the reaction rate exhibits the form of two square pulses with a minimal and maximal value $k_{basal}$ and $k_{max}$, respectively. The values of $k_{max}$ and the number of channels $n_{cha}$ were set to match an ATP consumption rate of $7 \cdot 10^5$ ATPs/s. The basal level of ATP consumption reflects additional housekeeping processes and is given in Table 4. For the synaptic simulations, we assumed a spontaneous firing rate of 5 Hz and vesicle release probability of 0.25 leading to a number of $\sim 1.8 \cdot 10^4$ ATP molecules per second needed at the synapse.

##### VDAC

In our model, we included VDACs to let ATP molecules exit the mitochondria into the cytosol. Since we were not aiming at capturing specific features of the channels, we implemented a rather simple Markov chain model that assumes that VDAC proteins interact with ATP and translocate it into the cytosol by the chemical reaction ($\text{VDAC} + \text{ATP}_{\text{mito}} \rightleftharpoons \text{VDAC} + \text{ATP}_{\text{cyto}}$). Based on experimental estimations of the VDAC density (De Pinto et al., 1987), we determined the number of channels $n_{vdac}$ as $1.1 \cdot 10^4$ for our specific mitochondrion model (Table 3). The rate constant of the reaction was set such that ATP export was not substantially delayed by the interaction with VDACs. Hence, in simulations with ATP molecules diffusing from a spherical region in the interior of the OM to the cytosol, the VDAC dependent export reduced the cytosolic ATP concentration to 75 % of the VDAC independent export within 10 milliseconds.

#### 2.2 Space independent ODE approach

For each of the model components, we derived an ODE system assuming mass action kinetics (Keener and Sneyd, 1998). The equations were integrated with PyDSTool (Clewley et al., 2007) using a Radau integrator.

##### ANT ODE model

The ODE system for the ANT model is given by Equations 1. The variables L, TL, LT, etc. represent the number of molecules in each state. One ANT can be found in one of the 11 states described in Figure 10. For the 11 variables, we derived 10 differential equations and used the conservation of total number of proteins

$L_{tot} = L + DL + DLD + DLD + TL + LT + TLT + TLT' + DLT + TLD$ from which we deduced the number of molecules in the state DLT. The governing equations read

$$
\begin{aligned}
\frac{dL}{dt} &= -(k^f_{T_o}\, T_o + k^f_{T_i}\, T_i + k^f_{D_o}\, D_o + k^f_{D_i}\, D_i)\, L + k^b_{T_o}\, TL + k^b_{T_i}\, LT + k^b_{D_o}\, DL + k^b_{D_i}\, LD \\
\frac{dTL}{dt} &= k^f_{T_o}\, T_o\, L - (k^b_{T_o} + k^f_{D_i}\, D_i + 2k^f_{T_i}\, T_i)\, TL + k^b_{T_i}\, TLT + k^b_{T_i}\, TLT^{'} + k^b_{D_i}\, TLD \\
\frac{dLT}{dt} &= k^f_{T_i}\, T_i\, L - (k^b_{T_i} + k^f_{D_o}\, D_o + 2k^f_{T_o}\, T_o)\, LT + k^b_{T_o}\, TLT + k^b_{T_o}\, TLT^{'} + k^b_{D_o}\, DLT \\
\frac{dDL}{dt} &= k^f_{D_o}\, D_o\, L - (k^b_{D_o} + k^f_{T_i}\, T_i + k^f_{D_i}\, D_i)\, DL + k^b_{D_i}\, DLD + k^b_{D_i}\, DLD^{'} + k^b_{T_i}\, DLT \\
\frac{dLD}{dt} &= k^f_{D_i}\, D_i\, L - (k^b_{D_i} + k^f_{T_o}\, T_o + k^f_{D_o}\, D_o)\, LD + k^b_{D_o}\, DLD + k^b_{D_o}\, DLD^{'} + k^b_{T_o}\, TLD \\
\frac{dTLD}{dt} &= k^f_{D_i}\, D_i\, TL + k^f_{T_o}\, T_o\, LD - (k^b_{D_i} + k_{cp})\, TLD + k_p\, DLT - k^b_{T_o}\, TLD \\
\frac{dDLD}{dt} &= k^f_{D_o}\, 0.5\, D_o\, LD + k^f_{D_i}\, 0.5\, D_i\, DL - (k^b_{D_o} + k^b_{D_i} + k_d)\, DLD + k_d\, DLD^{'} \\
\frac{dDLD^{'}}{dt} &= k^f_{D_o}\, 0.5\, D_o\, LD + k^f_{D_i}\, 0.5\, D_i\, DL - (k^b_{D_o} + k^b_{D_i} + k_d)\, DLD^{'} + k_d\, DLD \\
\frac{dTLT}{dt} &= k^f_{T_o}\, 0.5\, T_o\, LT + k^f_{T_i}\, 0.5\, T_i\, TL - (k^b_{T_o} + k^b_{T_i} + k_t)\, TLT + k_t\, TLT^{'} \\
\frac{dTLT^{'}}{dt} &= k^f_{T_o}\, 0.5\, T_o\, LT + k^f_{T_i}\, 0.5\, T_i\, TL - (k^b_{T_o} + k^b_{T_i} + k_t)\, TLT^{'} + k_t\, TLT
\end{aligned} \qquad (1)
$$

Here, $D_i$ and $T_i$ describe the number of ADP and ATP molecules in the matrix and $D_o$ and $T_o$ in the IMS and ICS building the outside space, respectively. $T_{cyto}$ describes the number of ATP molecules in the cytosol. The rate constants are labeled according to the reaction they drive, e.g. for reaction $T_o + L \rightarrow TL$, the corresponding rate constant is $k^f_{T_o}$ and for the inverse reaction $k^b_{T_o}$. The reactions that translocate metabolites are: DLT $\xrightarrow{k_p}$ TLD, exporting ATP to the outside and ADP to the inside, TLD $\xrightarrow{k_{cp}}$ DLT bringing ATP inside and ADP outside, TLT $\xrightarrow{k_t}$ TLT' transporting ATP from inside to outside and vice versa (this reaction matters for example when ATP within the media is labeled and we used this flux to determine the parameter $k_t$ from experimental data (Kraemer and Klingenberg, 1982)), and finally DLD $\xrightarrow{k_d}$ DLD' that analogously exchanges to ADP from the inside and outside. All parameters of the ANT model are summarized in Table 1.

##### ATP synthase ODE model

The ATP synthase model has six states represented in Figure 11A. In our simulations, n is set to 3 (i.e. 3 protons are needed to produce 1 molecule of ATP). As for the ANT, the total number of proteins is a conserved quantity ($E_{tot}$) leading to the conservation expression $E^{-3} + EH_3 + H_3E + H_3ES + H_3E^* + {}^{-3}E = E_{tot}$. Thus, we derived a system of 5 ODEs (Equations 2) to describe the ATP synthase dynamic by

$$
\begin{aligned}
\frac{dE^{-3}}{dt} &= -k_{65}\, E^{-3} + k_{56}\, EH_3 + k_{16}\, {}^{-3}E - k_{61}\, E^{-3} \\
\frac{d\, {}^{-3}E}{dt} &= -k_{16}\, {}^{-3}E + k_{61}\, E^{-3} - k_{12}\, {}^{-3}E + k_{21}\, H_3E \\
\frac{dH_3E^*}{dt} &= -k_{45}\, H_3E^* + k_{54}\, EH_3 + k_{34}\, H_3ES - k_{43}\, H_3E^*\, D_i \\
\frac{dH_3ES}{dt} &= k_{43}\, D_i\, H_3E^* - k_{34}\, H_3ES + k_{23}\, H_3E\, T_i - k_{32}\, H_3ES \\
\frac{dH_3E}{dt} &= -k_{23}\, H_3E\, T_i + k_{32}\, H_3ES - k_{25}\, H_3E + k_{52}\, EH_3 + k_{12}\, {}^{-3}E - k_{21}\, H_3E
\end{aligned} \qquad (2)
$$

and one more variable is deduced from the conservation expression. All parameter values are given in Table 2.

##### Metabolite ODE model

We also derived equations for the number of ATP and ADP molecules in the different compartments (Equations 3) given by

$$\begin{aligned}
\frac{dD_i}{dt} &= -(k^f_{D_i}\, L + k^f_{D_i}\, TL + k^f_{D_i}\, DL + k_{43}\, H_3 E_o)\, D_i + k^b_{D_i}\, LD + k^b_{D_i}\, TLD + k^b_{D_i}\, DLD + k^b_{D_i}\, DLD^{'} + k_{34}\, H_3 ES \\
\frac{dT_i}{dt} &= -(k^f_{T_i}\, L + k^f_{T_i}\, TL + k^f_{T_i}\, DL + k_{23}\, H_3 E)\, T_i + k^b_{T_i}\, LT + k^b_{T_i}\, DLT + k^b_{T_i}\, TLT + k^b_{T_i}\, TLT^{'} + k_{32}\, H_3 ES \\
\frac{dT_o}{dt} &= -(k^f_{T_o}\, L + k^f_{T_o}\, LT + k^f_{T_o}\, LD + k_p\, n_{vdac})\, T_o + k^b_{T_o}\, TL + k^b_{T_o}\, TLD + k^b_{T_o}\, TLT + k^b_{T_o}\, TLT^{'} + k_p\, T_{cyto}\, n_{vdac} \\
\frac{dT_{cyto}}{dt} &= n_{vdac}\, k\, T_o - n_{vdac}\, k\, T_{cyto} - n_{cha} k_{cha}(t) T_o \qquad (3)
\end{aligned}$$

For $\mathrm{T_{cyto}}$ the rate depends on the VDAC model parameters (Table 3) as well as on ATP consuming reactions. Therefore, when ATP consuming reactions were considered, the following term was added to the equation $-\mathrm{n_{cha} k_{cha}(t) \cdot T_o}$ with $\mathrm{k_{cha}(t)}$ being the rate of the reaction. This function has the form of two square pulses with $\mathrm{t_{on}}$ being the activation time and $\mathrm{t_{off}}$ the inactivation time for each pulse detailed in Table 4.

The rate constant of the bimolecular reactions have been normalized to have proper units to compute the number of particles, i.e. $\mathrm{k} = \frac{\mathrm{k'}}{\mathrm{N_a Vol}}$, where $\mathrm{N_a}$ is the Avogadro's number, and Vol represents the volume. Therefore, Vol is either the matrix volume or the volume of the outside space (ICS together with IMS).

#### 2.3 Estimation ATP production in synapses

Based on the turnover rate of ANTs and the number of translocators in the synaptic mitochondrion, we estimated the number of ATP molecules that can be exported to the cytosol per second (Table 5). The product of the turnover rate times the number of ANTs gives the maximum number of ATP molecules that can reach the cytosol per second if the ANTs are not the limiting step. The total amount of ANT proteins in synaptic mitochondria has been estimated to 1.37 nmol/mg protein (Chinopoulos et al., 2009) assuming 1 nmol/mg protein $\approx$ 1.25 mM (Magnus and Keizer, 1997) leading to a concentration of 1.71 mM. The inner membrane has a volume of 0.021 $\mu\mathrm{m}^3$ what correspond to $\sim 2 \cdot 10^4$ ANTs within the mitochondrion. This number of ANTs working at a turnover rate of 23 $\mathrm{s}^{-1}$ exports $4.6 \cdot 10^5$ ATPs/s. We further estimated the number of ATP produced by non-synaptic brain mitochondria (Table 5). This value critically depends on the volume of the mitochondria and, since no complete reconstructions are available in the literature, this is only a rough estimation based on an assumed cylindrical shaped mitochondria with a radius of 0.3 µm and length 1 µm (Perkins et al., 2001).

The number of ATP molecules produced by a synaptic mitochondrion coincides with the estimations of Attwell and Laughlin (Attwell and Laughlin, 2001) explained above. Moreover, a similar value of $6.02 \cdot 10^5$ ATP molecules per second has been also estimated for hair cell mitochondria (Babu et al., 2017).

#### 2.4 Time scale analysis

We calculate the spatial time scales for a diffusion coefficient of $\mathrm{D} = 1.5 \cdot 10^{-7}\mathrm{cm^2 s^{-1}}$ by considering the characteristic length scales as the diameter of cristae junctions $\sim$ 25.5 nm [2] ($\mathrm{L_1}$) and the distance from the center of the mitochondrion to the IBM $\sim$ 0.13 µm ($\mathrm{L_2}$). The time required to transverse these distances is $\tau$ and is calculated as $\tau = \mathrm{L^2/6D}$ leading to

$$\begin{aligned}
\tau_1 &= \frac{L_1^2}{6D} \sim 7 \cdot 10^{-6} s \\
\tau_2 &= \frac{L_2^2}{6D} \sim 2 \cdot 10^{-4} s\,.
\end{aligned}$$

A number of reactions occur in the ANT model. We calculated the corresponding time scales of some of the reactions to compare with the temporal scales of diffusion. The time scales of the forward reactions are denoted by $\tau_f$ and for the backward reactions by $\tau_b$ (Andrews and Arkin, 2006). For these calculations, we used the initial concentrations. As before L represents the free protein, $\mathrm{D_i}$ denotes ADP molecules from the matrix side and $\mathrm{D_o}$ ADP from the inner boundary membrane and cristae side (called outside) yielding

[2] The diameter of the cristae junctions is an average over 25 junctions measured in the reconstructed mesh.

$$D_i + L \rightleftharpoons LD \rightarrow \tau_f \sim 2.4 \cdot 10^{-4}s,\ \tau_b \sim 2.5 \cdot 10^{-5}s$$
$$D_o + L \rightleftharpoons DL \rightarrow \tau_f \sim 4 \cdot 10^{-4}s,\ \tau_b \sim 1 \cdot 10^{-2}s$$
$$T_i + L \rightleftharpoons LT \rightarrow \tau_f \sim 1.6 \cdot 10^{-3}s,\ \tau_b \sim 2.5 \cdot 10^{-5}s$$
$$T_o + L \rightleftharpoons TL \rightarrow \tau_f \sim 27s,\ \tau_b \sim 5 \cdot 10^{-3}s\,.$$

A similar analyzes has been done for the reactions of the ATP synthase model resulting in

$$E_o \rightleftharpoons EH_3 \rightarrow \tau_f \sim 1 \cdot 10^{-3}s,\ \tau_b \sim 1 \cdot 10^{-3}s$$
$$D_i + H3E_O \rightleftharpoons H3ES \rightarrow \tau_f \sim 3.1 \cdot 10^{-1}s,\ \tau_b \sim 1 \cdot 10^{-2}s$$
$$H3ES \rightleftharpoons T_i + H3E \rightarrow \tau_f \sim 2 \cdot 10^{-4}s,\ \tau_b \sim 1.8 \cdot 10^{-2}s\,.$$

Different time scales are found for the reactions of the ANT and ATP synthase models that spread over a large range but most of them are between $1 \cdot 10^{-5}$ s and $1 \cdot 10^{-2}$ s (Figure 12). We graphically represent the time scales of the reactions (blue dots) and the temporal scales of diffusion (black lines). They overlap for both considered diffusion coefficients but for the smaller diffusion coefficient the scales get closer together and diffusion limitation becomes more predominant.

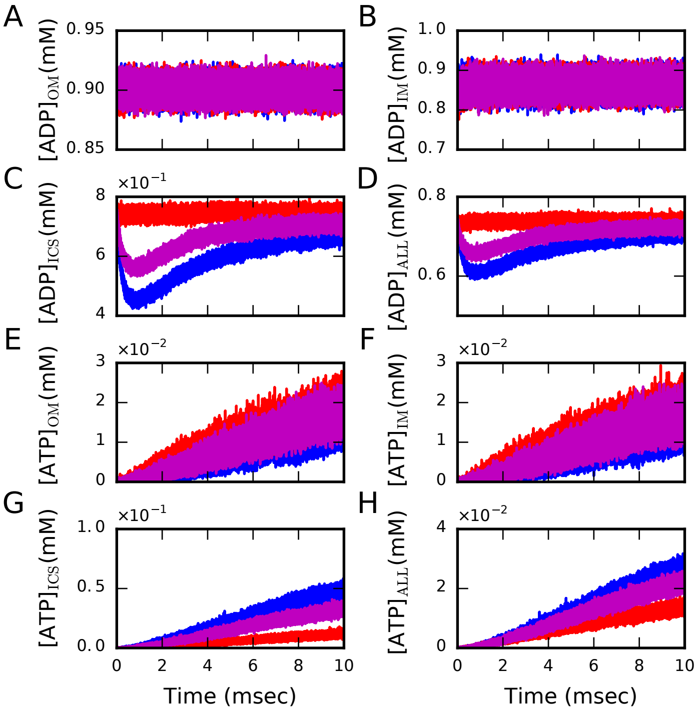

Figure 1: Local concentrations of the non-equilibrium dynamics of the synaptic mitochondrion driven by clamped ADP concentration and ATP export. We estimated the concentration of ADP (A-D) and ATP (E-H) in the cristae, inner boundary and outer membrane by counting the number of hits the corresponding surface experiences and approximating a small volume close to it. As before, ANT molecules were placed at three different locations: ANTs homogeneously distributed in the IBM (red); ANTs colocalized with ATP synthase at the most curved region of the CM (blue); ANTs in both locations (magenta).

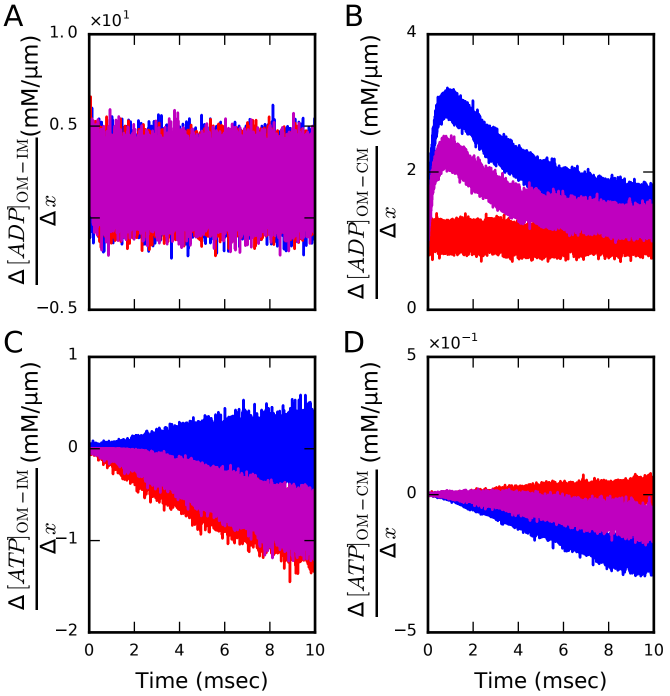

Figure 2: Local gradients of the non-equilibrium dynamics of the synaptic mitochondrion driven by clamped ADP concentration and ATP export. We estimated the concentration gradients of ADP (A-B) and ATP (C-D) formed between the inner boundary and outer membrane (A and C) as well as between the cristae and outer membrane (B and D). To calculate the gradients, estimated concentration were divided by the measured distances between the inner boundary and outer membrane (~0.02 μm) and between the middle of the mitochondrion to the outer membrane (~0.15 μm). Again ANT molecules were placed at three different locations: ANTs homogeneously distributed in the IBM (red); ANTs colocalized with ATP synthase at the most curved region of the CM (blue); ANTs in both locations (magenta).

| **Parameter** | **Value** | **Unit** |
|---|---|---|
| $k^{b}_{T_i}$ | $4 \cdot 10^4$ | $s^{-1}$ |
| $k'^{f}_{T_i}$ | $6.4 \cdot 10^6$ | $M^{-1}s^{-1}$ |
| $k^{b}_{T_o}$ | 200 | $s^{-1}$ |
| $k'^{f}_{T_o}$ | $4 \cdot 10^5$ | $M^{-1}s^{-1}$ |
| $k^{b}_{D_i}$ | $4 \cdot 10^4$ | $s^{-1}$ |
| $k'^{f}_{D_i}$ | $4 \cdot 10^6$ | $M^{-1}s^{-1}$ |
| $k^{b}_{D_o}$ | 100 | $s^{-1}$ |
| $k'^{f}_{D_o}$ | $4 \cdot 10^6$ | $M^{-1}s^{-1}$ |
| $k_p$ | 800 | $s^{-1}$ |
| $k_{cp}$ | 0.75 | $s^{-1}$ |
| $k_d$ | 0.48 | $s^{-1}$ |
| $k_t$ | 0.58 | $s^{-1}$ |
| Temperature | 310 | K |
| # ANT | $2 \cdot 10^4$ | |
| $\Delta\phi$ | -180 | mV |

Table 1: Chemical kinetic rate constants for the ATP/ADP translocator model.

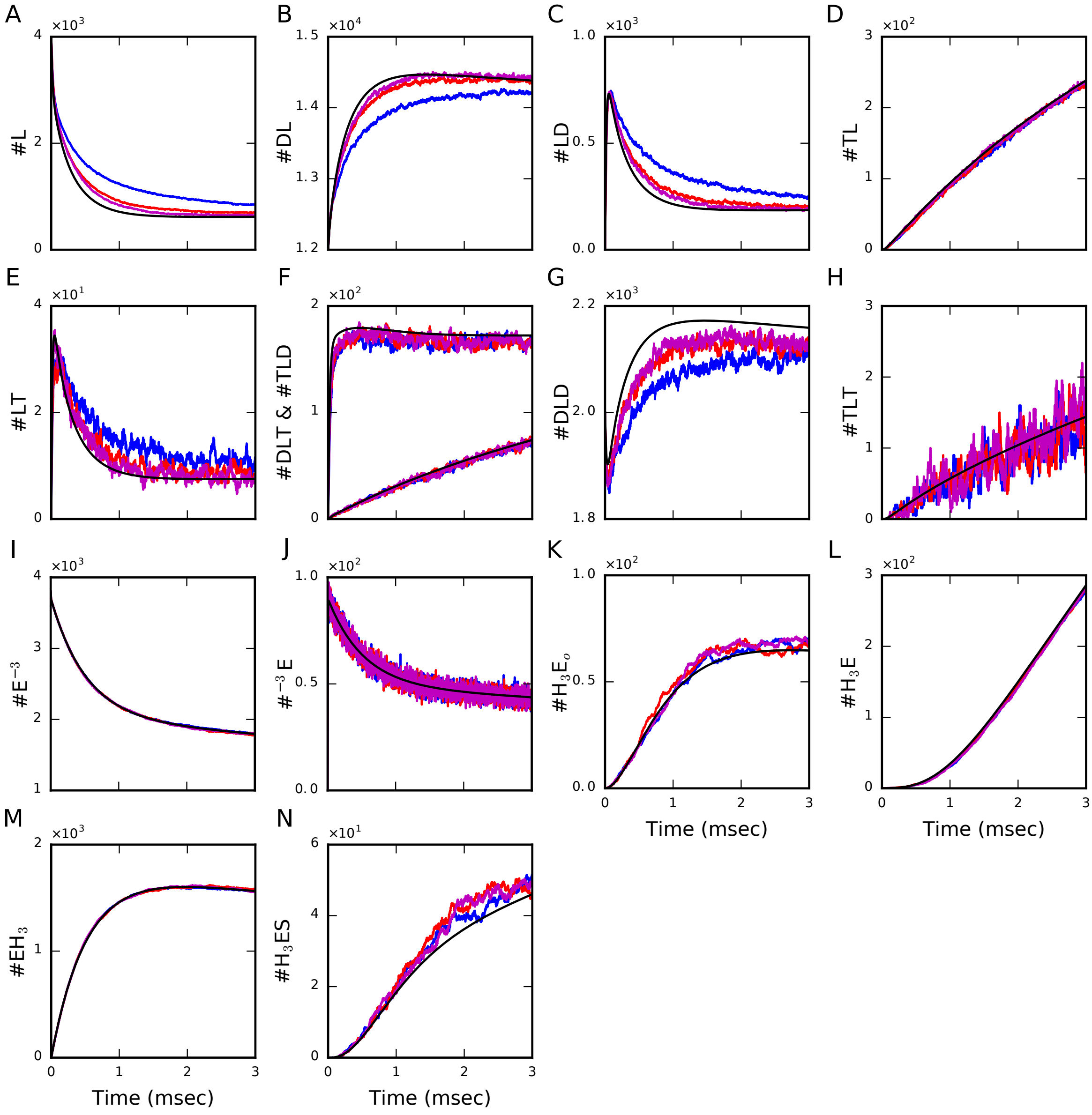

Figure 3: Additional variables for the isolated scenario of dynamic equilibration does not exhibit a strong dependence on the spatial arrangement. Comparison of molecule numbers between averaged trajectories of 10 different initial conditions in MCell and the results obtained with the ODE system. Simulations started with a a large number of ADP molecules outside (IMS and ICS) that were subsequently phosphorylated. ATP molecules accumulate outside due to the lack of VDACs. ANT molecules were placed in three different locations: ANTs homogeneously distributed in the IBM (red), ANTs colocalized with ATP synthase at the most curved region of the CM (blue) and ANTs in both locations (magenta). The results of the ODE system are shown in black. Panels A-H show the variables of the ANT model and panels I-N the variables of the ATP synthase model. The different arrangements of ANTs do not exhibit significant differences with the corresponding ODE system.

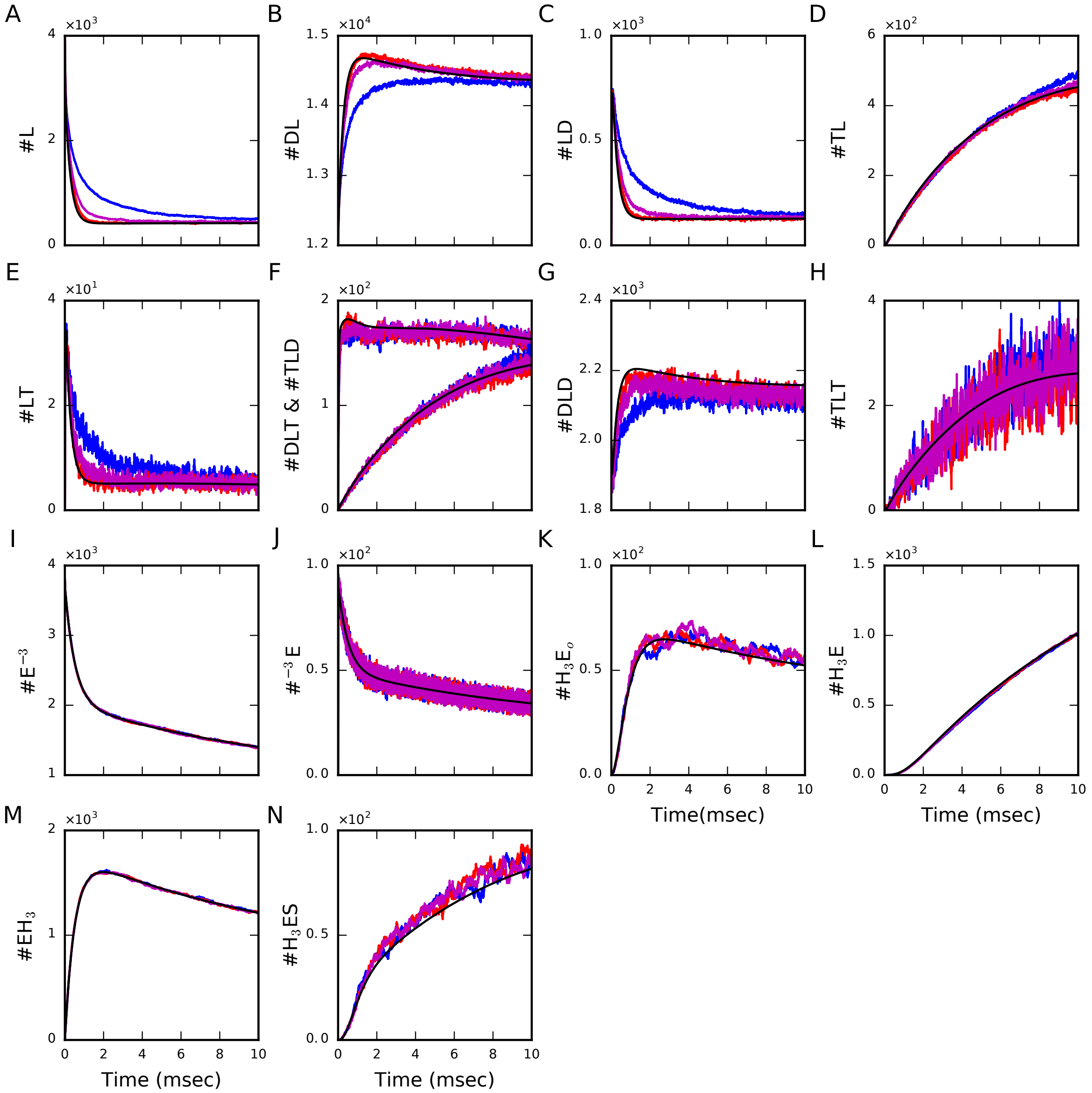

Figure 4: Additional variables for the non-equilibrium dynamics of the synaptic mitochondrion driven by clamped ADP concentration and ATP export. Averaged molecule number trajectories for the distinct ANT localizations with ANTs homogeneously distributed in the IBM (red), colocalized with ATP synthase at the most curved region of the CM (blue) or in both locations (magenta) were compared with results of the ODE system. This comparison exhibits most significant spatial effects for colocalization (blue). In panels A-H, we show variables of the ANT model and in panels I-N variables of the ATP synthase model.

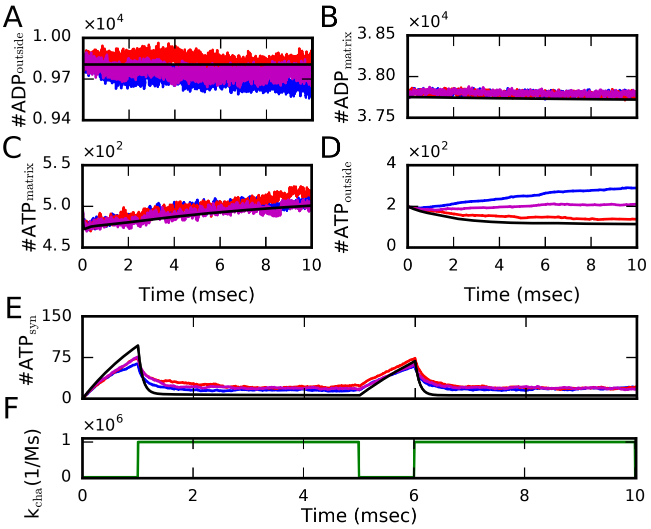

Figure 5: The energy dynamics at a presynaptic terminal. (*A*) In this configuration, the mitochondrion is placed within the segmented synapse (grey surface) and we considered ATP-consuming reactions. (*B*-*F*) Comparison of averaged molecule trajectories in the different compartments (color) and corresponding ODE results (black) for simulations of an action potential arrival at the presynaptic terminal by modulating the rate constant $k_{\text{cha}}$ of the ATP-consuming reactions (*G*). As before, the concentration of ADP in the OM was clamped and VDAC channels included in the OM. ANT molecules were placed in three different locations: ANTs homogeneously distributed in the IBM (red), ANTs colocalized with ATP synthase at the most curved region of the CM (blue), and ANTs in both locations (magenta).

| **Parameter** | **Value** | **Units** | **Reference** |
|---|---:|---:|---:|
| $\text{pH}_{\text{matrix}}$ | 7.8 | | (Song et al., 2013) |
| $\text{pH}_{\text{cristae}}$ | 7.4 | | (Song et al., 2013) |
| $[P_i]$ | 20 | mM | (Magnus and Keizer, 1997) |
| $\Delta\phi$ | -180 | mV | |
| Density | 2500 | $\mu\text{m}^{-2}$ | (Song et al., 2013; Schwerzmann et al., 1986) |
| # ATP synthase | 3800 | | |
| Temperature | 310 | K | |
| $k_{16}$ | 452457 | $s^{-1}$ | (Pietrobon and Caplan, 1985) |
| $k_{65}$ | $1 \cdot 10^3$ | $s^{-1}$ | (Pietrobon and Caplan, 1985) |
| $k_{54}$ | 100 | $s^{-1}$ | (Pietrobon and Caplan, 1985) |
| $k'_{43}$ | $8 \cdot 10^5$ | $M^{-1}s^{-1}$ | (Magnus and Keizer, 1997) |
| $k_{32}$ | $5 \cdot 10^3$ | $s^{-1}$ | (Pietrobon and Caplan, 1985) |
| $k_{21}$ | 40 | $s^{-1}$ | (Pietrobon and Caplan, 1985) |
| $k_{61}$ | 11006 | $s^{-1}$ | (Pietrobon and Caplan, 1985) |
| $k_{12}$ | 24 | $s^{-1}$ | (Pietrobon and Caplan, 1985) |
| $k'_{23}$ | 4 | $\mu M^{-1}s^{-1}$ | (Magnus and Keizer, 1997) |
| $k_{34}$ | 100 | $s^{-1}$ | (Pietrobon and Caplan, 1985) |
| $k_{45}$ | 100 | $s^{-1}$ | (Pietrobon and Caplan, 1985) |
| $k_{56}$ | 1000 | $s^{-1}$ | (Pietrobon and Caplan, 1985) |
| $k_{25}$ | $1.17 \cdot 10^{-12}$ | $s^{-1}$ | (Magnus and Keizer, 1997) |
| $k_{52}$ | 2 | $s^{-1}$ | (Magnus and Keizer, 1997) |
| n | 3 | | (Pietrobon and Caplan, 1985) |

Table 2: Chemical kinetic rate constants and parameters for the ATP synthase model.

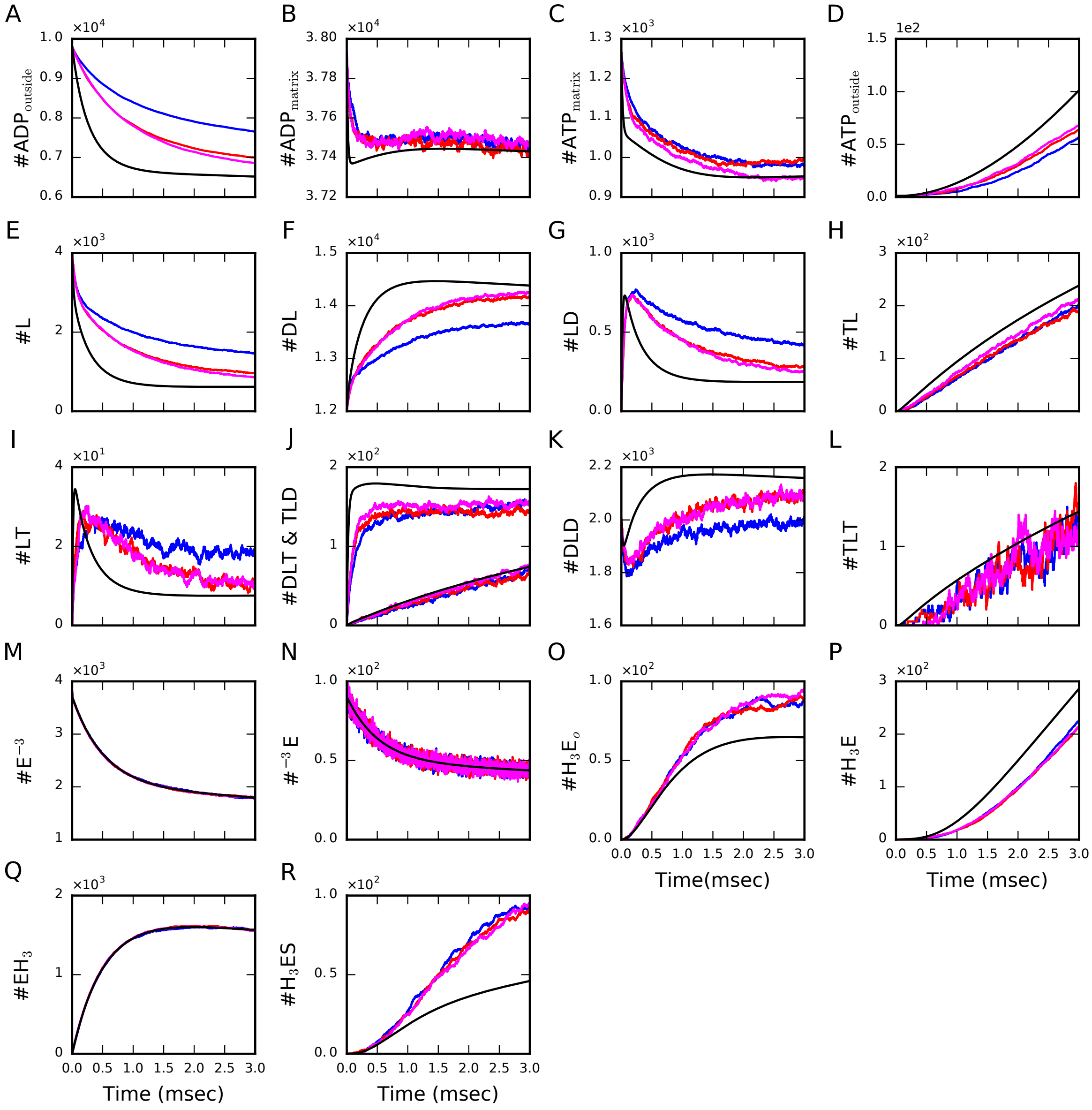

Figure 6: Concentration dynamics and variables for the isolated scenario of dynamic equilibration with reduced diffusion. ANTs molecules were again placed in three different locations: ANTs homogeneously distributed in the IBM (red), ANTs colocalized with ATP synthase at the most curved region of the CM (blue) and ANTs in both locations (magenta). Panels A-L show variables of the ANT model and panels M-R the variables of the ATP synthase model.

| Parameter | Value | Unit |
|---|---|---|
| $k'$ | $1 \cdot 10^6$ | $M^{-1}s^{-1}$ |
| Density | $1 \cdot 10^4$ | $\mu m^{-2}$ |
| $n_{vdac}$ | $7.1 \cdot 10^3$ | |

Table 3: Chemical kinetic rate constants for the VDAC model.

| Parameter | Value | Unit |
|---|---|---|
| $n_{cha}$ | $7 \cdot 10^5$ | |
| $k'_{basal}$ | $2.5 \cdot 10^4$ | $M^{-1}s^{-1}$ |
| $k'_{max}$ | $1 \cdot 10^6$ | $M^{-1}s^{-1}$ |
| $t^1_{on}$ | 1 | ms |
| $t^1_{off}$ | 5 | ms |
| $t^2_{on}$ | 6 | ms |
| $t^2_{off}$ | 10 | ms |

Table 4: Chemical kinetic rate constants for the ATP consuming reactions.

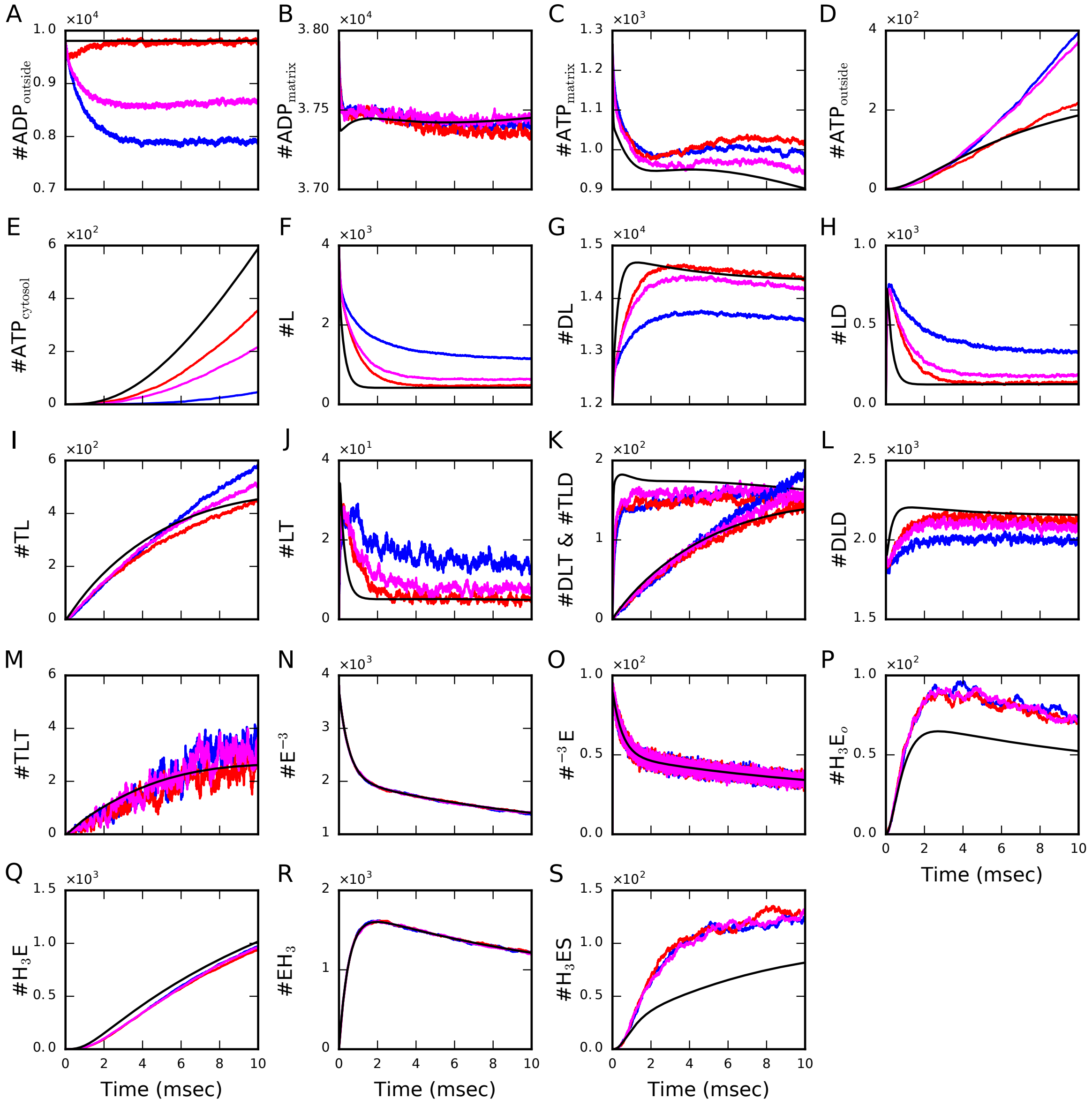

Figure 7: Concentration dynamics and variables for the non-equilibrium dynamics of the synaptic mitochondrion driven by clamped ADP concentration and ATP export with reduced diffusion. Comparison of averaged molecule number trajectories for the distinct ANT localizations (ANTs homogeneously distributed in the IBM (red), ANTs colocalized with ATP synthase at the most curved region of the CM (blue) and ANTs in both locations (magenta)) with results of the ODE system (black) exhibits a most significant spatial effects for colocalization (blue). Panels (A-L) show variables of the ANT model and panels M-R variables of the ATP synthase model.

| **Tissue** | **Rate**($s^{-1}$) | **Density**(nmol/mg prot) | **Concentration** (mM) | **Volume** ($\mu m^3$) | **Nr.ANTs** | **ATPs/sec** | **ATP moles/sec** |
|---|---|---|---|---|---|---|---|
| Synapse | 23 $^\star$ | 1.37$^\star$ | 1.71 ‡ | 0.021 $^\dagger$ | $2 \cdot 10^4$ | $4.6 \cdot 10^5$ | $0.75 \cdot 10^{-18}$ |
| Non-synapse | 22 $^\star$ | 1.44 $^\star$ | 1.79 ‡ | 0.28 | $3 \cdot 10^5$ | $6.6 \cdot 10^6$ | $1.1 \cdot 10^{-17}$ |

Table 5: Estimation ATP production in brain tissue.

$^\star$ Chinopoulos et al. (2009), ‡ 1 nmol/mg prot ≈ 1.25 mM (Magnus and Keizer, 1997), $^\dagger$ our estimation.

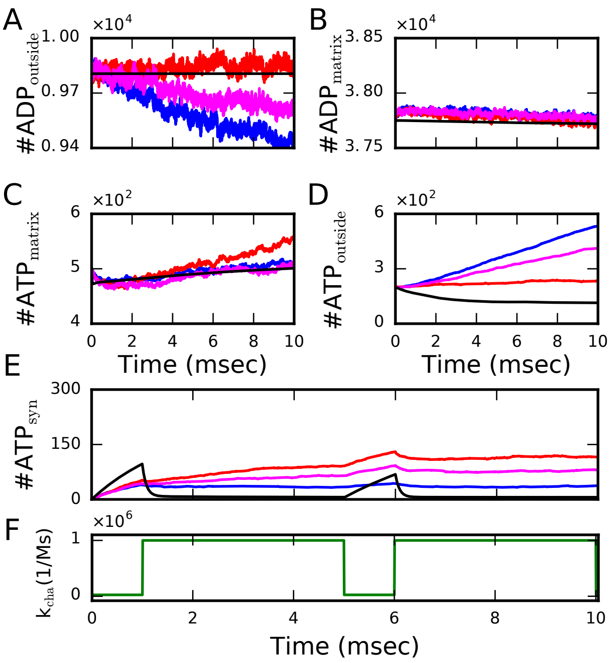

Figure 8: The energy dynamics of a presynaptic terminal with reduced diffusion compared to Figure 3 in the main text. Comparison of averaged trajectories of the number of molecules in different compartments with the results obtained by the ODE system. In this configuration, the mitochondrion is placed in its physiological context, the presynaptic terminal, and ATP consuming reactions were considered to mimic the arrival of an action potential at the terminal. For this purpose, we emulated the increase of the rate constant of the ATP consuming reactions (F). As before, the concentration of ADP in the OM was clamped and VDACs were included in the OM. ANT molecules were placed in three different locations: ANTs homogeneously distributed in the IBM (in red), ANTs colocalized with ATP synthase at the most curved region of the CM (blue) and ANTs in both locations (magenta). The results of the ODE system are shown in black.

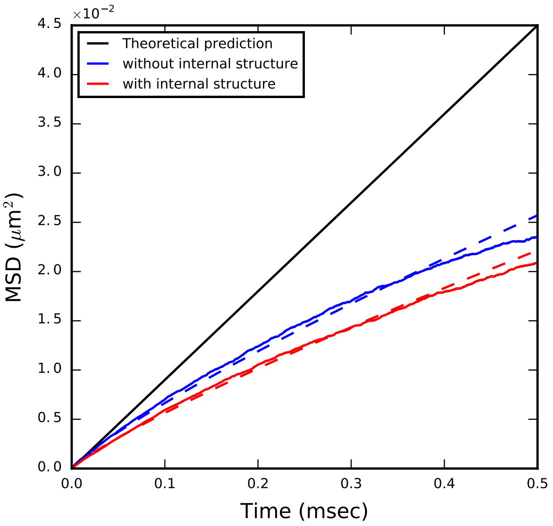

Figure 9: Time courses of the mean square displacement (MSD) for particles diffusing in the matrix without considering the internal structure (blue) and with consideration of the internal structure (red). Initially, 2500 particles were distributed homogeneously in the matrix, and their position was monitored to compute the MSD for both configurations. The dash lines represent a power law fitting of the data. The black line is the theoretical prediction of the MSD for Fick's law of diffusion.

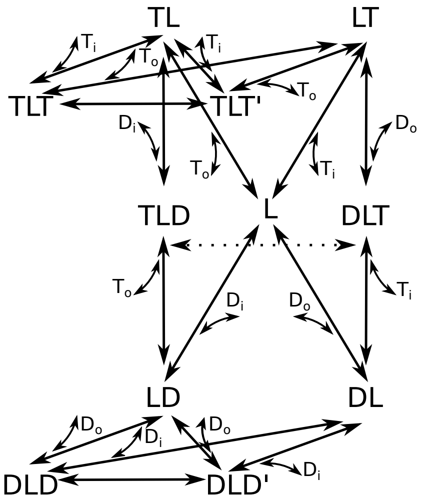

Figure 10: Markov chain model of ANT describing the molecular kinetics. ATP and ADP are represented by the letter T and D, respectively, and index i refers to the matrix side and o to the IMS and ICS together (called outside). L represents the free protein and YLX a triple molecular state with one X molecule bound from the matrix side and one Y molecule bound from the outside.

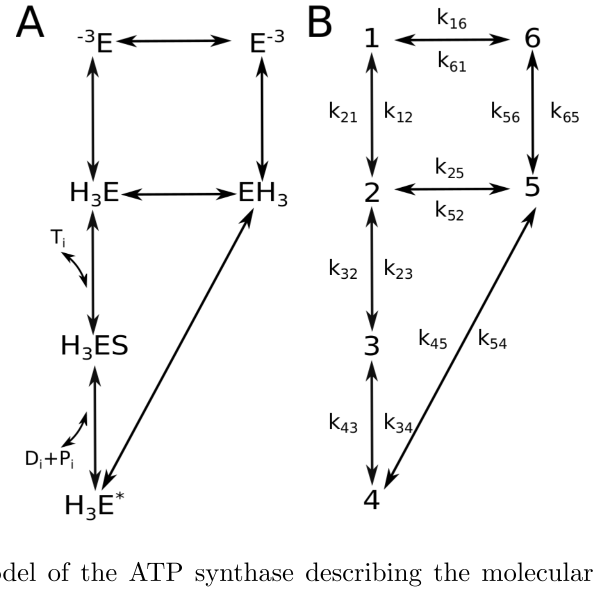

Figure 11: Markov chain model of the ATP synthase describing the molecular kinetics. The ATP synthase model is composed of six states. (A) States toward the matrix are on the left and oriented toward the outside are on the right. A clockwise cycle starting from $E^{-n}$ corresponds to the binding of n protons from the outside, their transport to the matrix and subsequent binding of ADP and $P_i$ to the synthesis of ATP, and unbinding of the n protons in the matrix. (B) The productive cycle with corresponding state numbering and state transition rate constants $k_{ij}$ describing transitions from state i to state j.

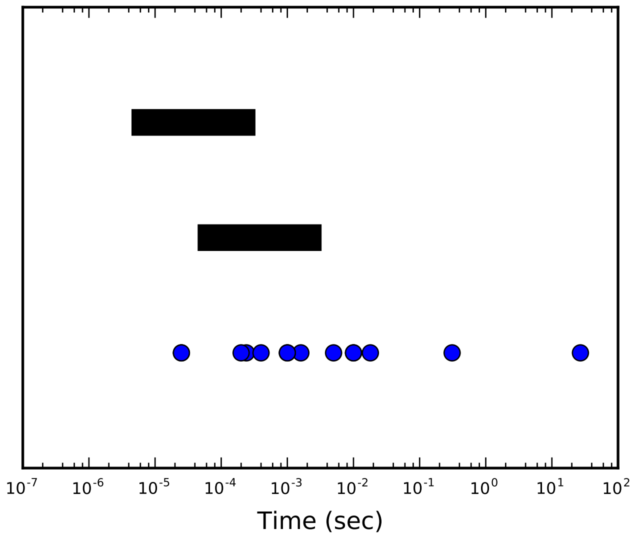

Figure 12: Graphical representation of the temporal scales. The black lines represent the temporal scales of diffusion with the upper line holds for a diffusion coefficient of $D = 1.5 \cdot 10^{-7} \mathrm{cm}^2\mathrm{s}^{-1}$ and the lower line for a diffusion coefficient of $D = 1.5 \cdot 10^{-8} \mathrm{cm}^2\mathrm{s}^{-1}$. The blue dots are a representation of the temporal scales of the reactions. Decreasing the diffusion coefficient brings the temporal scales closer.
